# ELQ-331 as a prototype for extremely durable chemoprotection against malaria

**DOI:** 10.1101/687756

**Authors:** Martin J. Smilkstein, Sovitj Pou, Alina Krollenbrock, Lisa A. Bleyle, Rozalia A. Dodean, Lisa Frueh, David J. Hinrichs, Yuexin Li, Thomas Martinson, Myrna Y. Munar, Rolf W. Winter, Igor Bruzual, Samantha Whiteside, Aaron Nilsen, Dennis R. Koop, Jane X. Kelly, Stefan H. I. Kappe, Brandon K. Wilder, Michael K. Riscoe

## Abstract

**Background:** The potential benefits of long-acting injectable chemoprotection (LAI-C) against malaria have been recently recognized, prompting a call for suitable candidate drugs to help meet this need. On the basis of its known pharmacodynamic and pharmacokinetic profiles after oral dosing, ELQ-331, a prodrug of the parasite mitochondrial electron transport inhibitor ELQ-300, was selected for study of pharmacokinetics and efficacy as LAI-C in mice.

**Methods:** Four trials were conducted in which mice were injected with a single intramuscular dose of ELQ-331 or other ELQ-300 prodrugs in sesame oil with 1.2% benzyl alcohol; the ELQ-300 content of the doses ranged from 2.5 to 30 mg/kg. Initial blood stage challenges with *Plasmodium yoelii* were used to establish the model, but the definitive study measure of efficacy was outcome after sporozoite challenge with a luciferase-expressing *P.yoelii*, assessed by whole-body live animal imaging. Snapshot determinations of plasma ELQ-300 concentration ([ELQ-300]) were made after all prodrug injections; after the highest dose of ELQ-331 (equivalent to 30 mg/kg ELQ-300), both [ELQ-331] and [ELQ-300] were measured at a series of timepoints from 6 hours to 5 ½ months after injection.

**Results:** A single intramuscular injection of ELQ-331 outperformed four other ELQ-300 prodrugs and, at a dose equivalent to 30 mg/kg ELQ-300, protected mice against challenge with *P. yoelii* sporozoites for at least 4 ½ months. Pharmacokinetic evaluation revealed rapid and essentially complete conversion of ELQ-331 to ELQ-300, a rapidly achieved (< 6 hours) and sustained (4-5 months) effective plasma ELQ-300 concentration, maximum ELQ-300 concentrations far below the estimated threshold for toxicity, and a distinctive ELQ-300 concentration vs. time profile. Pharmacokinetic modeling indicates a high-capacity, slow-exchange tissue compartment which serves to accumulate and then slowly redistribute ELQ-300 into blood, and this property facilitates an extremely long period during which ELQ-300 concentration is sustained above a minimum fully-protective threshold (60-80 nM).

**Conclusions:** Extrapolation of these results to humans clearly predicts that ELQ-331 should be capable of meeting and far-exceeding currently published duration-of-effect goals for antimalarial LAI-C. Allometric scaling from mice to humans would predict a several-fold enhancement in the relationship between duration-of-effect and dose, and available drug engineering and formulation technologies would be expected to offer significant improvement over the simple powder in sesame oil used here. Furthermore, the distinctive pharmacokinetic profile of ELQ-300 after treatment with ELQ-331 may facilitate durable protection using a variety of delivery and formulation options, and may enable protection for far longer than 3 months. Particularly in light of the favorable pharmacodynamic profile of ELQ-300, ELQ-331 warrants consideration as a leading prototype for LAI-C.

## Background

Despite a remarkable recent reduction in the global burden of malaria, there remains an urgent need for novel antimalarial drug treatments. It is evident that new drugs are needed to maintain efficacy in the face of emerging drug resistance, but even in the absence of drug resistance there are settings and sub-populations in which drugs are unavailable, unaffordable, impractical, ineffective, or unsafe. These concerns clearly apply to current prevention and treatment applications and are only magnified when future eradication efforts are also considered. For some needs, many antimalarial drug options exist, for others only a few, and for some none at all. For this reason, thought-leaders in the field have sought to identify critical gaps in antimalarial drug options, to define the ideal characteristics of drugs to fill each gap, and to disseminate that information in the form of Target Product Profiles (TPPs) (1).

Until recently, TPPs have focused on maintaining or improving the effectiveness of drugs for conventional uses: treatment of malaria, prophylaxis in the non-immune facing new malaria exposure, and intermittent prophylaxis in critical sub-groups among populations with endemic exposure. Missing from this list has been any role for drugs to achieve sustained, large scale malaria prevention among populations in endemic regions. It has been long-hoped that a vaccine could provide the effective and durable protection needed for that purpose, and long-assumed that drug treatment could not feasibly achieve it. The importance and the urgency of finding options to provide effective and durable protection has increased as strategies to eradicate malaria have been considered. Whether by protecting the whole population early in the process, or more likely by targeting critical groups late in the process, it is evident that sustained, reliable protection against infection and transmission will be an absolute requirement for future successful eradication efforts.

The results of vaccine studies to-date indicate that a feasible, highly-effective, and durably-protective vaccine will not be available soon (2). At the same time, an increasing body of information has demonstrated the safety and efficacy of long-acting injectable (LAI) medications to treat other conditions (3-6), and pilot studies have shown durable protection against malaria in mice after intramuscular (IM) drug injection (7, 8). Together, these observations prompted the Medicines for Malaria Venture (MMV) to develop the first drug TPP for long-acting injectable chemoprotection (LAI-C) against malaria (9).

As a starting point, the MMV suggested the following priorities for LAI-C candidates: (1) Multi-stage activity including liver and blood stage schizonticidal, and transmission-blocking efficacy, (2) rapid-onset high efficacy, sustained for 3 months, (3) minimal or no consequential drug-drug interactions, (4) high safety and tolerability both at the injection site and systemically, including in pediatric patients and those with glucose-6-phosphate dehydrogenase (G-6-PD) deficiency, and (5) formulations with suitable viscosity (injectable through small-bore needles), shelf-life (>2-3 years in tropical conditions), and low cost. The history of drug resistance development during blood stage infection has led to the recommendation that two drugs with the above features be combined for LAI-C, but it is acknowledged that a single drug might prove adequate, given the likely diminished risk of de novo drug resistance during causal prophylaxis. Because drugs used for LAI-C will be present at the injection site, in blood, and in tissues for a very long time period, safety testing will be more complex, lengthy, and costly than that for other drug applications. Since any prior toxicology information will therefore be essential to down-select appropriate drugs for LAI-C, the MMV has suggested dividing candidates for LAI-C into three conceptual groups: tier 1, comprised of existing pharmaceuticals with available clinical safety information; tier 2, new drugs which can be delivered orally for initial safety testing; and tier 3, drugs that can only be tested parenterally because adequate oral delivery is not possible.

Very few antimalarials provide causal prophylactic efficacy, so tier 1 candidates include only atovaquone, with or without proguanil; the anti-folates, sulfadoxine and pyrimethamine; and the 8-aminoquinolines, primaquine and tafenoquine. While each has attractive features, each has liabilities. Atovaquone has a remarkable propensity to rapidly generate high-grade resistance (10), although that risk is likely less during causal prophylaxis and may be further mitigated by the diminished transmissibility of highly atovaquone-resistant parasites (11). To match a partner drug and provide effective coverage for three months, the relatively short elimination half-life of proguanil (12) would mandate a very large delivered dose and attendant possible toxicity risk, or extensive formulation engineering to control release from the injection site. The antifolates face widespread, pre-existing drug resistance (13), and although antifolate-caused life-threatening dermatologic toxicity (e.g., Stevens-Johnson syndrome) is uncommon (14), the risk likely precludes consideration of these drugs for LAI-C. Although recent studies indicate the safety and efficacy of some 8-aminoquinoline dosages in some patients with G-6-PD deficiency (15), LAI-C would result in months of continuous oxidative stress making it almost certain that G-6-PD testing would be required prior to use, and likely that G-6-PD deficiency would be a contraindication for 8-aminoquinoline LAI-C. Furthermore, 8-aminoquinolines could be ineffective in the small but significant proportion of the population lacking adequate CYP 2D6 to form active metabolites (16). From among these tier 1 candidates, atovaquone may be the most promising, but no single drug or two-drug combination appears likely to provide sufficiently feasible, effective, and safe LAI-C in the context of eradication efforts in regions of high endemicity.

ELQ-300 (see Table 1) is an experimental antimalarial with causal prophylactic, blood stage schizonticidal, and transmission-blocking activity (17). It acts at the Q_i_ site of cytochrome bc_1_ to selectively inhibit parasite mitochondrial electron transport and inter-related pyrimidine biosynthesis (17, 18), thus preventing nucleic acid synthesis. ELQ-300 has low nanomolar efficacy in vitro against blood stage *Plasmodium falciparum, vivax (17)*, and *knowlesi* (personal communication, D.A. van Schalkwyk) including multidrug-resistant laboratory strains and clinical isolates, and distinctly notable transmission-blocking potency (20). Compared to atovaquone, which acts at the Q_o_ site of cytochrome b and rapidly gives rise to drug resistance in vitro, the propensity to develop ELQ-300 resistance is far lower (>10^−5^ for atovaquone, <10^−8^ for ELQ-300) (17, 21). Inhibition of cytochrome bc_1_ electron transport by ELQ-300 is also far more parasite-specific (IC_50_ *P. falciparum* = 0.56 nM, IC_50_ human = >10,000 nM) when compared to atovaquone (IC_50_ *P. falciparum* = 2 nM, IC_50_ human = 460 nM) (17). In extensive pre-clinical testing, ELQ-300 has an excellent safety profile, no evident in vitro drug-drug interactions, and slow elimination which facilitates single-dose cures in mice. Oral bioavailability is adequate to prove efficacy and to evaluate pharmacokinetics (PK) in animals, but the poor aqueous solubility and high crystallinity of ELQ-300 limits the extent and predictability of exposure after oral dosing, making it unsuitable as an orally-delivered drug, despite its other favorable features.

**Table 1:**
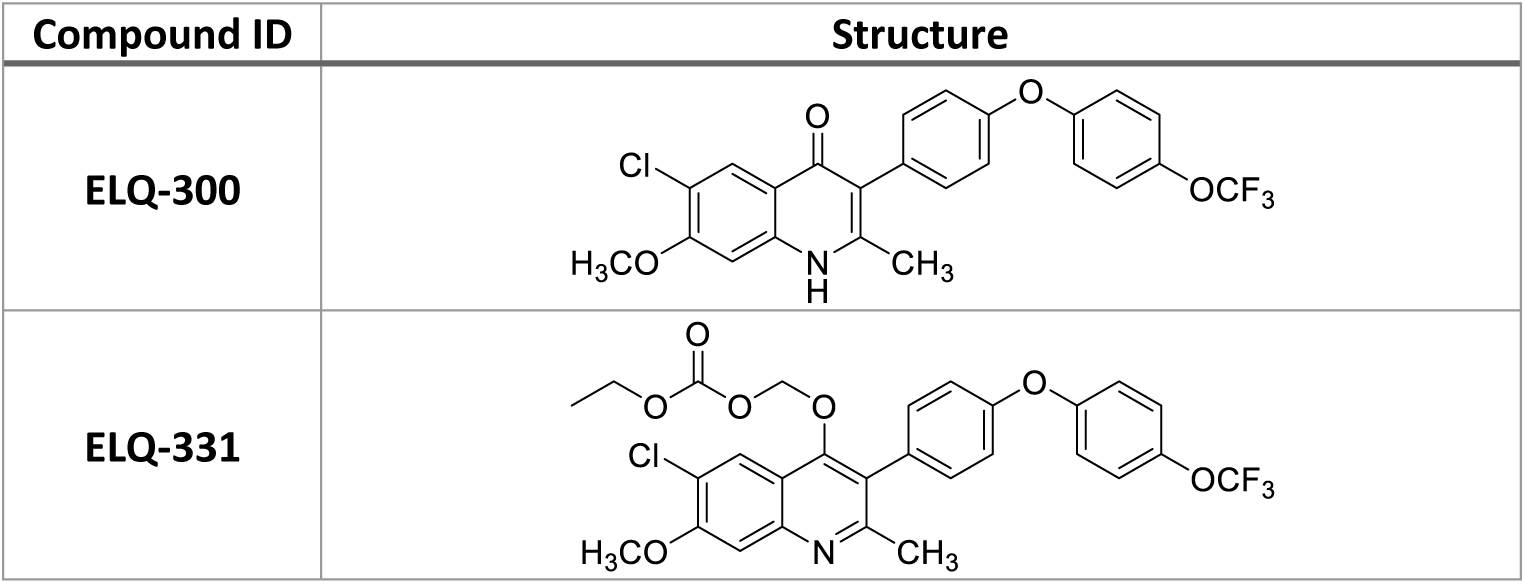
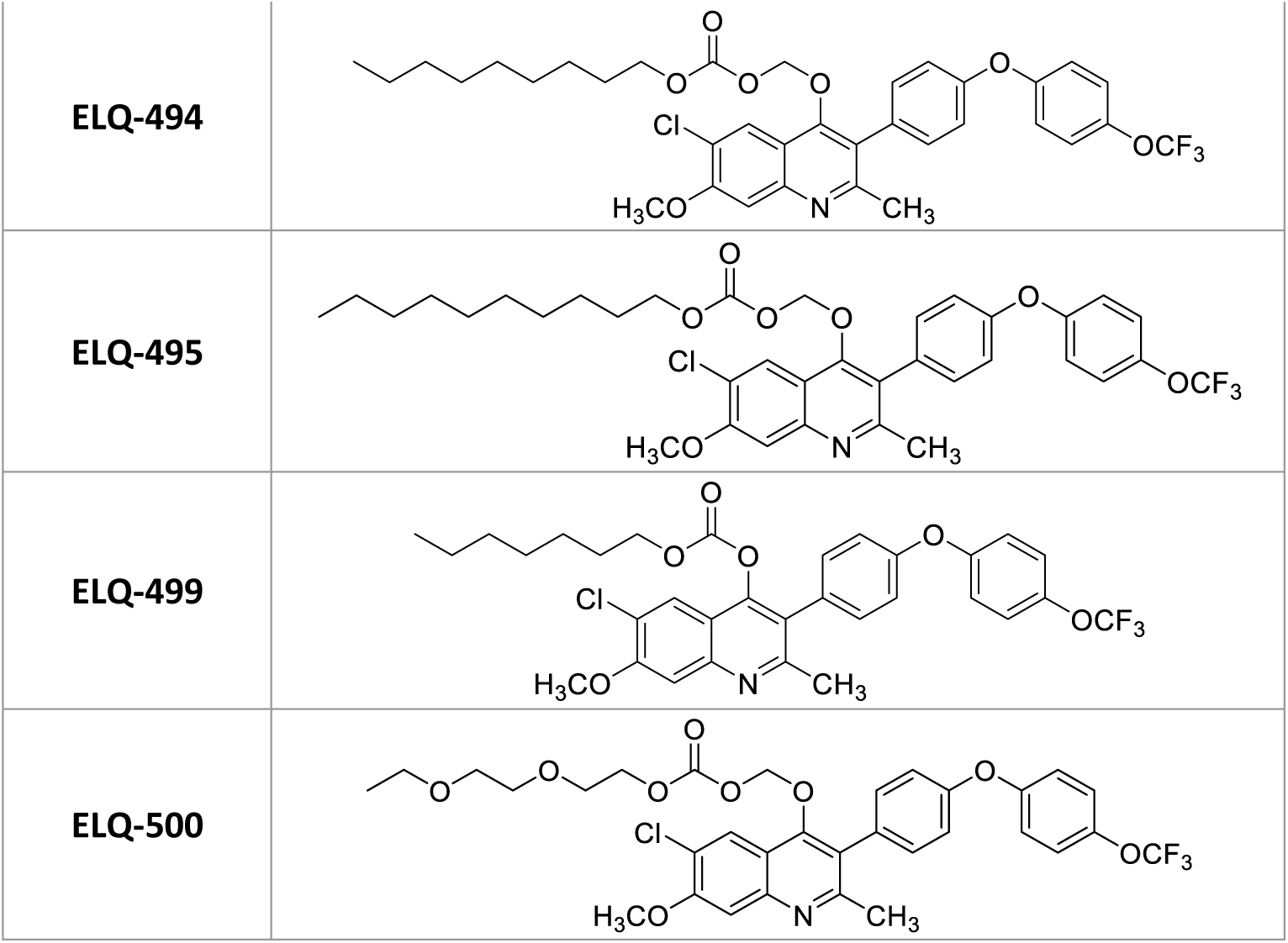
Structures of ELQ-300, ELQ-331, and other ELQ-300 prodrugs included in this report : The synthesis, chemical analysis, physicochemical properties, and other details related to ELQ-300 and ELQ-331 are previously published (17, 19). ELQ-494, ELQ-495, ELQ-499, and ELQ-500 will be described as part of a future report covering the entire scope of the ELQ-300 prodrug project.

| Compound ID | Structure |
| --- | --- |
| ELQ-300 |  |
| ELQ-331 |  |
| **ELQ-494** |  |
| **ELQ-495** |  |
| **ELQ-499** |  |
| **ELQ-500** |  |

To improve oral bioavailability while preserving the highly desirable features of ELQ-300, we have sought to develop prodrugs of ELQ-300 that are better absorbed from the gastrointestinal tract but then are rapidly converted to ELQ-300 (19, 22). ELQ-331 (see Table 1), an alkoxycarbonate ester of ELQ-300, has emerged as a leading contender. ELQ-331 also has limited aqueous solubility, but with far less tendency to form insoluble crystals and, probably on that basis, results in much-increased ELQ-300 exposure after oral dosing. In mice, the in vivo efficacy of ELQ-331 by gavage is superior to ELQ-300 (19), and in rats, 10 mg/kg of an ELQ-331 formulation resulted in a mean maximum plasma [ELQ-300] of 12.6 μM (manuscript submitted). ELQ-331 has undergone successful efficacy testing against blood stage infection and sporozoite challenge in mice (19), and toxicology assessment rats (see Additional File 1); and we continue to advance ELQ-331 and other prodrugs for oral use. It is evident, however, that the pharmacodynamic and PK features of ELQ-331 are ideal for a LAI-C candidate, and that physicochemical features which are problematic for oral dosing might be beneficial for its use as a LAI-C. We therefore undertook the following studies in mice to assess the efficacy and PK profile of IM ELQ-331 in sesame oil, and to conduct a limited comparison with other ELQ-300 prodrugs.

## Methods

### ELQ-300 prodrug treatment in mice

All treatments were administered in 0.05 mL of sesame oil containing 1.2% benzyl alcohol (v/v), injected through a 0.625-inch, 25 gauge needle into the caudal thigh of mice weighing 25-35 gm. To facilitate comparisons, prodrug doses were calculated and are expressed on the basis of ELQ-300 content. For example, mice in “10 mg/kg” ELQ-331 treatment groups received 12.1 mg/kg of ELQ-331, the molar equivalent of 10 mg/kg of ELQ-300. Stable solubility of each prodrug in the injection vehicle was determined prior to dosing (see Additional File 2).

### Blood stage challenges to assess activity against blood stage infection

Blood stage challenges were done using a lethal *P. yoelii*_*K*_ parasite (subsp. *yoelii*, Strain K (Kenya), BEI Resources, NIAID, NIH: MRA-428, contributed by Wallace Peters and Brian L. Robinson). Mice were injected intravenously via tail vein with 3 × 10^5^ *P. yoelii*_*K*_-infected erythrocytes and evaluated by microscopy of Giemsa-stained blood smears 5 days later. Mice that remained parasite-free were reassessed by blood smears weekly for at least 28 days, at which time they were scored as “not infected”. Mice with any evident parasitemia at any time point during the study were scored as “infected” and euthanized.

### Sporozoite challenges to assess causal prophylactic activity

Sporozoite challenges were done using a transgenic, non-lethal *P. yoelii* XNL parasite that constitutively expresses a GFP-luciferase fusion protein throughout the life cycle (Py-GFP-luc) (23). Female six to eight-week-old Swiss Webster (SW) mice were injected with Py-GFP-luc-infected blood to begin the growth cycle. The infected mice were used to feed female *Anopheles stephensi* mosquitoes after gametocyte exflagellation was observed. Ten days after the blood meal, 15–20 mosquitoes were dissected to evaluate midgut oocyst formation. At day 14 or 15 post-blood meal, salivary glands were removed, sporozoites liberated using a pestle and then isolated by centrifugation through glass wool. Sporozoites were then counted by hemocytometer and diluted with incomplete RPMI to a concentration of 5 × 10^5^ per mL. Samples were maintained on ice from the time of salivary gland dissection until final dilution in RPMI. Mice were then injected with 0.2 mL (10^5^ sporozoites) intravenously via tail vein and evaluated by whole-animal bioluminescence imaging 2 days after inoculation to assess liver infection, and 5-6 days after inoculation to determine if blood stage infection occurred.

Whole-animal imaging was done using the IVIS Spectrum CT and Living Image software (Perkin-Elmer). Mice were injected intraperitoneally with 150 mg/kg D-luciferin (XenoLight, Perkin-Elmer) in PBS, 10 minutes prior to imaging, then shortly before and throughout imaging, anesthesia was induced and maintained with isoflurane. If no luminescence signal was evident using auto exposure settings, exposure time was increased in stepwise fashion to at least 2 minutes at widest aperture. Optimal settings often varied between imaging sessions, but for all groups of mice compared head-to-head at any given time point, all camera and software settings were identical. Image assessment was qualitative only, to determine the presence or absence of evident infection.

### Blood sampling and processing for determination of plasma ELQ-331 and ELQ-300 concentration

Mice were warmed for 5-10 minutes under a lamp prior to sampling. A fifty microliters blood sample was collected from a tail vein using a calibrated, heparinized microcapillary tube, and transferred to a microfuge tube containing 5.7 μL of 500 mM sodium fluoride, maintained in an ice bath before and after sampling. This protocol had been previously shown by us to limit ex vivo esterase metabolism of ELQ-331 to <2% (see Additional File 2). At each time point, after all required mice were sampled, the specimens were moved to a cold room, centrifuged at high speed for 1 minute and then 20 μL of supernatant plasma was transferred to a new tube, which was immediately frozen at −80° C until analysis.

### Analytical methods for determination of plasma ELQ-331 and ELQ-300 concentration

Prodrug and drug concentrations were analyzed using an Applied Biosystems QTRAP 4000 (Foster City, CA) with electrospray ionization in positive mode. The mass spectrometer was interfaced to a Shimadzu (Columbia, MD) SIL-20AC XR auto-sampler followed by two LC-20AD XR LC pumps. The instrument was operated with the following settings: source voltage 5500 kV, GS1 20, GS2 50, CUR 15, TEM 600, and CAD MEDIUM. The scheduled Multiple Reaction Monitoring (MRM) transitions monitored are listed in the table below; the bold are used for quantification. Compounds were individually infused and instrument parameters optimized for each MRM transition. A gradient mobile phase was delivered at a flow rate of 0.3 mL/min; it consisted of two solvents: A - 0.1% formic acid in water and B - 0.1% formic acid in acetonitrile. Separation was achieved using a Dacapo DX-C18 column at 40°C and the autosampler was kept at 30°C to prevent the precipitation of ELQ and related compounds. The gradient elution was as follows: 40% B, held for 0.5 min; increase to 95% B at 3 min; 100% B was held for 2.5 min; decrease to 40% B over 0.1 min; hold at 10% B for 1.4 minutes to re-equilibrate. Method modification was necessary to separate ELQ-331 from the ELQ-300 as there was in-source fragmentation which initially gave erroneous values for ELQ-300 in the presence of ELQ-331. The retention times were 3.09 minutes for ELQ-300, 2.91 minutes for ELQ-316 (internal standard for ELQ studies [IS]) and 2.98 minutes for ELQ-331; separate curves were prepared routinely for ELQ-300 and ELQ-331. Data were acquired using Analyst 1.5.1 software and analyzed using Multiquant 3.0.2.

| Compound | RetentionTime<br>(min) | Q1<br>Mass | Q3<br>Mass | DP | EP | CE | CXP |
| --- | --- | --- | --- | --- | --- | --- | --- |
| **ELQ-300** | **3.09** | **476** | **433.1** | **166** | **10** | **61** | **12** |
| ELQ-300 | 3.09 | 476 | 426.1 | 166 | 10 | 61 | 12 |
| **ELQ-316**<br>**(IS)** | **2.91** | **460** | **417.1** | **176** | **10** | **59** | **12** |
| ELQ-316<br>(IS) | 2.91 | 460 | 239.1 | 176 | 10 | 83 | 16 |
| **ELQ-331** | **2.98** | **578** | **476** | **100** | **10** | **45** | **14** |
| ELQ-331 | 2.98 | 578 | 433.1 | 100 | 10 | 73 | 12 |

Plasma samples were quickly thawed at room temperature and crashed with freshly prepared 1:9 dimethylformamide:acetonitrile with 100 ng/mL ELQ-316 internal standard at a ratio of 5 μL plasma to 95 μL of crash solution. Samples were vortexed for 2 minutes on a pulsing vortexer, centrifuged for 2 minutes at high speed, and the supernatant transferred to sample vials. Prior to analysis the samples were pre-heated in the autosampler for a minimum of 45 minutes to ensure stable signal. A standard curve was prepared by spiking commercial mouse plasma with analyte in a range from 1 ng/mL to 2000 ng/mL, crashed and processed identically to test samples.

### Pharmacokinetic analysis

Group statistics (mean ± SEM) for plasma drug concentration values and graphical representation of concentration vs. time curves were done using Prism 7 software (GraphPad, San Diego). The apparent maximum concentration (C_max_) and the time from dosing to C_max_ (T_max_) were determined by inspection of the results table and graph. When apparent half-life (T_1/2_) is described for any curve segment, it was calculated by the formula: T_1/2_ = 0.693/k_el_, where k_el_ = - slope of the ln(concentration) vs time curve. Non-compartmental modeling and initial compartmental modeling was done using Phoenix WinNonLin 6.4, build 8 (Certara L.P., Princeton, NJ). Since WinNonLin lacks a built-in 3-compartment model with first-order absorption, final compartmental modeling was performed with SimBiology tools in MATLAB software (MathWorks, Natwock, MA). Output PK parameter estimates are considered to represent “apparent” rather than physiologic values.

### Summary of all trials

Four separate IM ELQ-331 treatment trials were ultimately performed, the rationale for which is described (see Results, below). For reference, Table 2 summarizes the various trials to be discussed. Female Swiss-Webster mice (CFW, Charles River Laboratories. Inc. Raleigh, NC) were used for Trials 1-3, and female CF-1 mice (Charles River Laboratories. Inc. Raleigh, NC) were used in Trial 4. On arrival, the animals were acclimated for at least 7 days in quarantine. The animals were housed in a cage maintained in a room with a temperature range of 64-79 °F, 34-68% relative humidity and a 12-h light/dark cycle. Food and water were provided *ad libitum* during quarantine and throughout the study. The animals were fed a standard rodent maintenance diet. The study design did not include a formal drug toxicity assessment, however, blood sampling afforded frequent opportunities for observation of the mice in addition to the standard, ongoing monitoring of animal welfare per protocols. Transient or ongoing abnormalities in animal behavior or health, atypical of the normal course of events during healthy animal housing, were required to be reported; none were noted.

**Table 2:** Summary of the four treatment trials described in this report

| Trial | Dose, prodrug | Time from dosing to testing |  |  |
| --- | --- | --- | --- | --- |
|  |  | Plasma [ELQ-300] determination | Blood stage challenge | Sporozoite challenge |
| 1 | 30 mg/kg ELQ-331 | 47, 60, 90, 120 days | 7, 14, 21, 40 days | 137 days |
|  | 30 mg/kg ELQ-499 |  |  |  |
|  | 30 mg/kg ELQ-500 |  |  |  |
| 2 | 30 mg/kg ELQ-331 | 6, 12, 18, 24, 36, 48, 72 hours;<br>weekly 1-10, 14, 18, 22, 24 weeks | n/a | 48 days |
| 3 | 10 mg/kg ELQ-331 | 30, 70, 77 days | n/a | 77 days |
|  | 10 mg/kg ELQ-494 | 30, 70 days |  | n/a |
|  | 10 mg/kg ELQ-495 | 30, 70 days |  |  |
|  | 10 mg/kg ELQ-501 | 30, 83 days |  |  |
| 4 | 2.5 mg/kg ELQ-331 | 14, 27 days | n/a | n/a |
|  | 5 mg/kg ELQ-331 |  |  | 29 days |
|  | 10 mg/kg ELQ-331 |  |  |  |

## Results

### Pilot testing for activity against blood stage infection and initial ELQ-300 snapshot PK

In Trial 1 (see Figure 1), mice were injected with 30 mg/kg of ELQ-331, ELQ-499, or ELQ-500 on Day 0; controls received no drug. To establish initial duration-of-effect parameters to guide subsequent studies, control mice and mice from each treatment group were challenged with *P. yoelii*_*K*_ -infected erythrocytes on Day 7, 14, 21, or 40 after treatment, using a new sub-group of mice (N=3-4) at each timepoint. Blood smears of all control mice, at all timepoints, showed patent infection after 5 days (mean parasitemia = 24 ± 4.9%). All treated mice remained parasite-free for at least 28 days.

**Figure 1:**
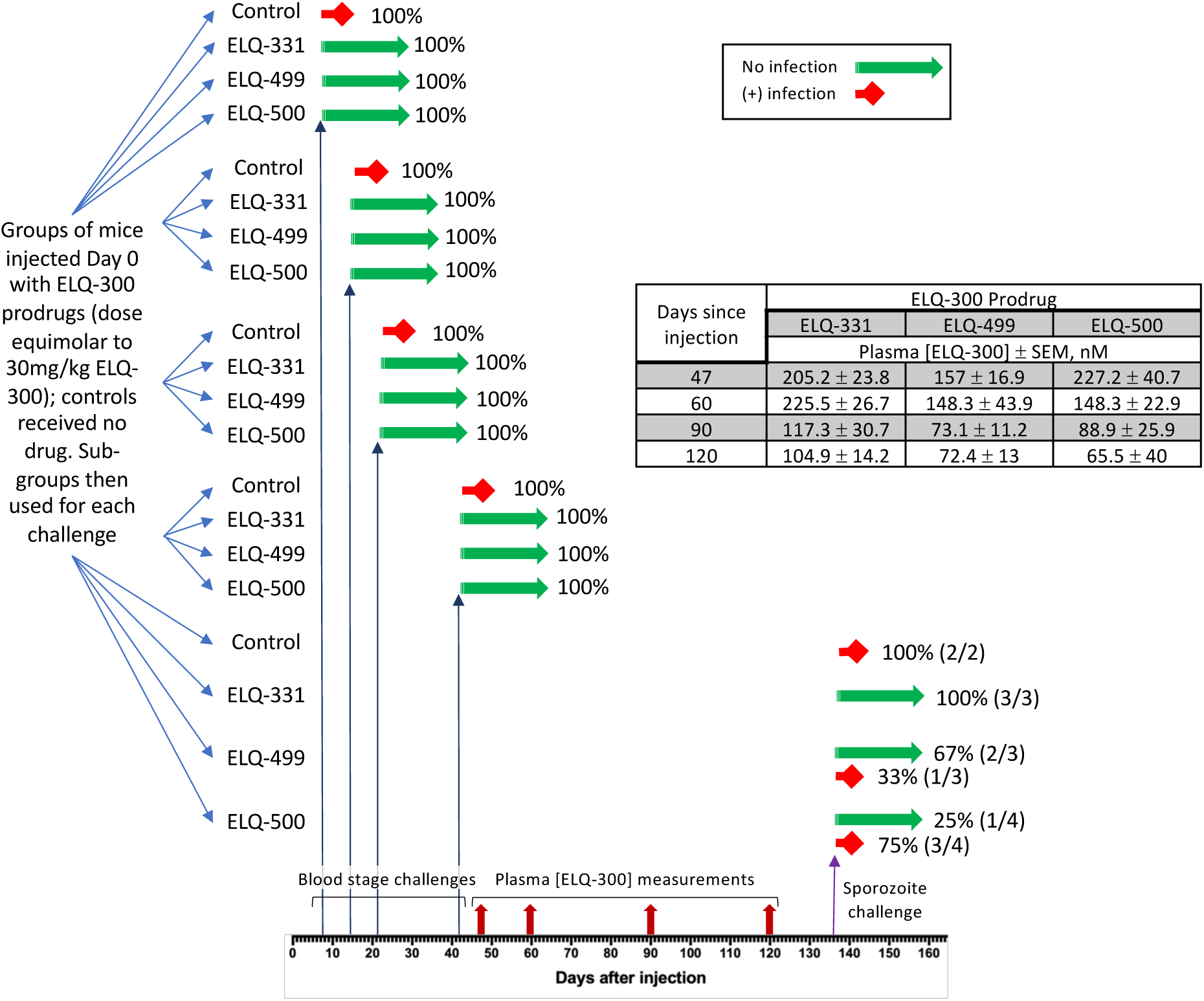
Summary of Trial 1 protocol and results The timepoints for blood stage challenges, blood sampling for snapshot pharmacokinetic analysis, and sporozoite challenges are noted on the x-axis. The outcomes of efficacy trials are represented graphically as red = no protection and green = complete protection, and the proportion of challenged mice with each outcome is noted. The table insert summarizes the results of snapshot PK at the indicated timepoints.

Since these efficacy results indicated the persistence of elevated blood ELQ-300 concentration ([ELQ-300]), we sought to quantify [ELQ-300] and compare values in the three prodrug treatment groups. Tail vein blood samples were obtained 47, 60, 90, and 120 days after treatment (see Figure 1, table insert). ELQ-331 produced a more sustained elevation of [ELQ-300] than the other tested prodrugs, remaining above 100 nM for 4 months. ELQ-331 was therefore selected for Trial 2, a more detailed PK evaluation (see Pharmacokinetics of 30 mg/kg ELQ-331, below), and the still-persistent elevation of [ELQ-300] prompted us to initiate sporozoite challenge testing in the remaining Trial 1 mice (see Causal prophylactic efficacy, below).

### Pharmacokinetics of 30 mg/kg IM ELQ-331 in sesame oil

In Trial 2, 24 mice were injected intramuscularly with 30 mg/kg ELQ-331 and then divided into 6 groups. Blood samples were obtained from one group at each timepoint in a repeating, sequential rotation, so that the same mice were sampled at every sixth timepoint. Samples were taken 6, 12, 18, 24, 36, 48, and 72 hours after injection, then weekly from 1 to 10 weeks, then every 4 weeks until 22 weeks, and a final sample was taken 24 weeks after dosing. Plasma [ELQ-331] and [ELQ-300] were measured at each time point (see Figure 2).

**Figure 2:**
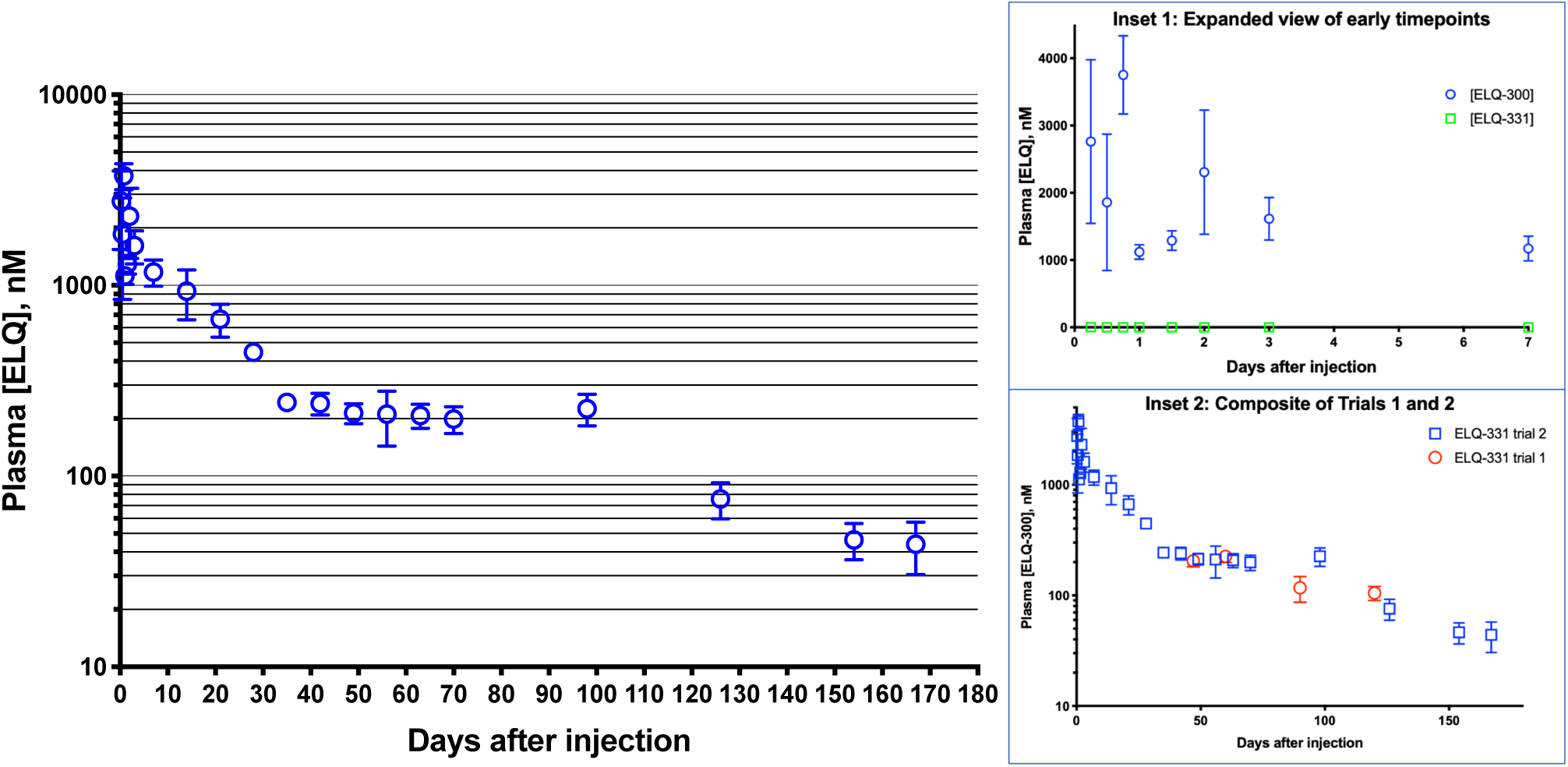
Pharmacokinetics of ELQ-300 after IM injection of 30 mg/kg ELQ-331 in Trial 2 Main figure summarizes Trial 2 PK results in a log-linear plot of [ELQ-300] (mean ± SEM) vs time. Inset 1 shows detail of days 0-7 in an expanded linear-linear plot of [ELQ-331] and [ELQ-300] vs time. Inset 2 shows a composite log-linear plot combining values from Trial 1 and Trial 2.

The lower limit of quantitation for [ELQ-331] was 1.2 nM, and values were below that cut-off or non-detectable in 86% of samples, even in the first days after treatment (see Figure 2, Inset 1). When quantifiable, [ELQ-331] levels were trivial (mean = 4 nM, max = 11.5 nM), never reaching 1% of the corresponding [ELQ-300]. Importantly, this indicates rapid and highly-efficient metabolism to ELQ-300.

Variability is evident in [ELQ-300] values at several time points within the first few days (see Figure 2, Inset 1). Remembering that each of the first 6 time points represents a different group of mice, we speculate that time points with very little variability (e.g., 24 and 36 hours) may represent consistent IM delivery to all mice in that group, whereas high variability within a group may indicate that the portion of the dose successfully deposited and retained within the muscle varied between mice. If true, then early time points with lower and more consistent values may provide a more accurate picture of early PK values after IM injection. Some of the highest early [ELQ-300] values may then represent the effect of a portion of the dose being deposited or extruded outside the muscle to a site where rapid access to the circulation can occur. Given the very small blood volume of a mouse, even a small fraction of the injected dose rapidly reaching the circulation could produce transient high μM [ELQ-300], so it is difficult to judge whether or not the early variability actually reflects any substantial variation in the proportion of the dose deposited intramuscularly. Our interpretation of [ELQ-300] values determined at later time points (see below) is that the early variability here does not indicate substantial loss of the dose from the injection site, and that successful dosing was broadly consistent. Since the objective is to rapidly reach effective but sub-toxic concentrations, it is notable that at every early timepoint, [ELQ-300] is well above 100 nM and does not approach concentrations associated with toxicity established in prior studies in rats (see Discussion). With the above caveat in mind, i.e., that several early observed timepoints may not be predictive of true IM PK, the ELQ-300 apparent C_max_ was 3752 ± 582 nM, at T_max_ = 18 hours.

The remaining concentration vs. time curve can be divided into 3 apparent phases. Between approximately 1 week and 1 month after injection, despite the early variability discussed above, groups show remarkably little variability and follow a consistent down-sloping curve with an apparent T_1/2_ = 14 days, far longer than the elimination T_1/2_ seen after intravenous dosing in mice (15 hours) (17). The long apparent T_1/2_ during this phase is a strong indication that absorption of ELQ-300 from the injection site into blood is ongoing. More speculatively, the consistency of PK results between and within groups during this phase also suggests that, despite the wide variability seen earlier in the same mice, a high proportion of the dose was consistently deposited intramuscularly. The next phase of the curve shows a decrease in the slope of the concentration vs time curve, which remains relatively flat between 1 and 3 months after injection. Inserting [ELQ-300] values from Trial 1 reveals remarkable overlap (see Figure 2, Inset 2) supporting the validity of the results. After day 90, the downward slope increases for the final phase of the curve with an apparent terminal T_1/2_ = 29.5 days.

A full discussion of the PK modeling analysis (dataset definition, models comparison, model diagram, expressions and equation) is presented in Additional File 3, the results of which will inform future analysis and simulations. The main finding relevant to the consideration of ELQ-331 as a prototype for LAI-C is the qualitative finding of a multi-compartment PK profile for ELQ-300, including a large-capacity, slow-exchange tissue compartment with significant impact on the duration of ELQ-300 exposure. This is demonstrated in Figure 3, showing the curve, parameter estimates, and analysis of fit resulting from fitting the “Edited” dataset using a 2-compartment model with first-order absorption and elimination (For dataset definition, model diagram, expressions and equation, see Additional File 3). Results confirm an excellent fit, with little difference between observed and predicted [ELQ-300] values, and evenly distributed residuals. Since the slope of the [ELQ-300] vs. time curve after 30 days cannot be attributed to either substantial ongoing absorption of ELQ-300 from the injection site (24, 25) or slow intrinsic elimination of ELQ-300 (17), it must be predominantly related to release from the large-capacity (Peripheral_vol_ = 4.89 L vs. Central_vol_ = 0.12 L), slow-exchange (intercompartmental clearance [Q12] = 0.075 L/h) compartment.

**Figure 3:**
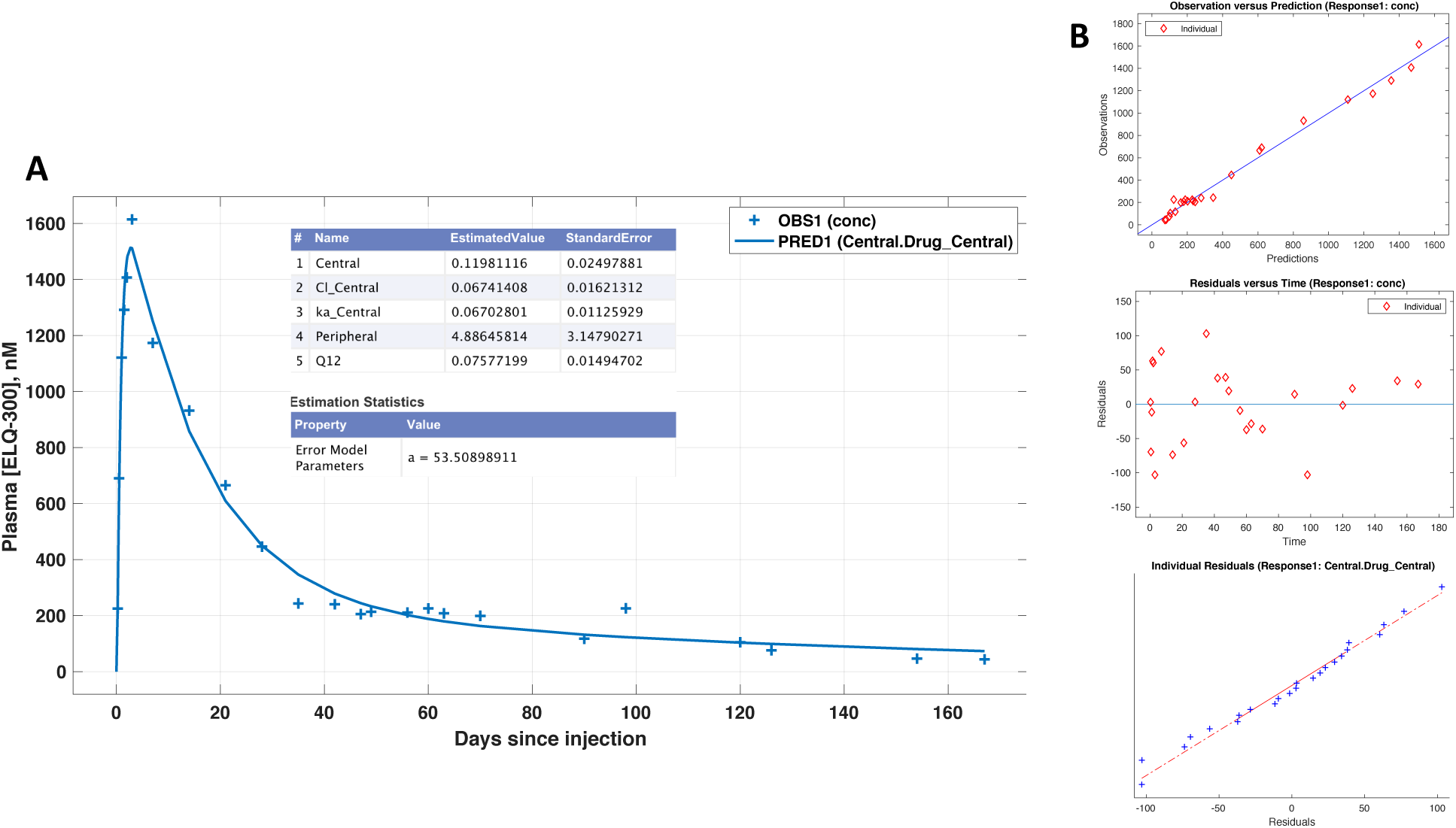
Results of 2-compartment model fit (A) Linear-linear plot of observed mean [ELQ-300] values at each timepoint (+ symbols) and values predicted from the curve fit (line). Estimated model parameters are shown in the insert table. Central = Apparent volume of the central compartment (L); Cl_Central = central compartment clearance (L/hr); ka_Central = rate constant for absorption; Peripheral = apparent volume of the tissue compartment; Q12 = intercompartmental clearance (L/hr). (B) Indicators of model fit validity: correspondence of observed and predicted values, unbiased distribution of residuals.

### Causal prophylactic efficacy of ELQ-331 after sporozoite challenge, corresponding plasma [ELQ-300], and estimation of MEC_LAI-C_

Mice treated with 30 mg/kg ELQ-331 from Trial 1 (137 days after treatment, N=3) and Trial 2 (48 days after treatment, N=4), and untreated controls (N=2) were included in initial Py-GFP-luc sporozoite challenge testing (see Table 3, upper segment). None of the treated mice developed infection; controls had both liver stage and subsequent blood stage infection evident by luminescence imaging. The finding that a protective [ELQ-300] was sustained for more than 4 months led us to initiate Trial 3, evaluating a lower dose (10 mg/kg) of ELQ-331 in a limited comparison with three additional ELQ-300 prodrugs.

**Table 3:**
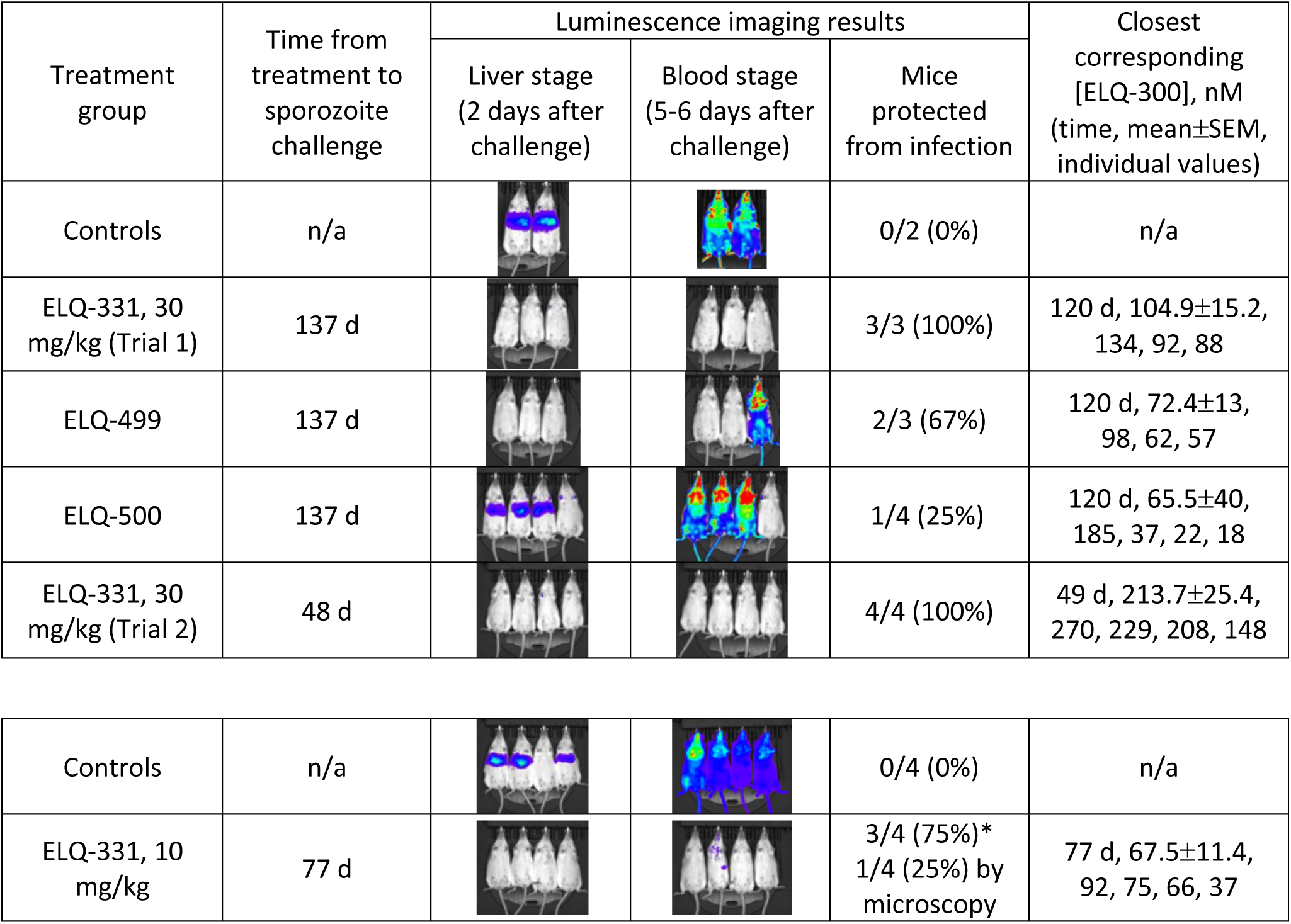
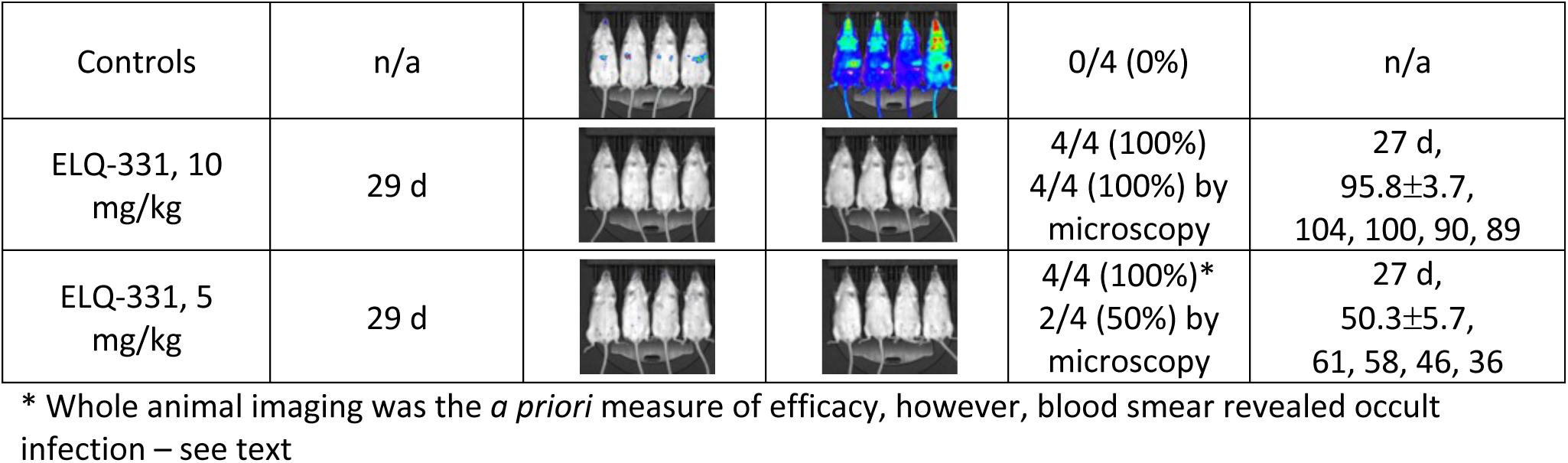
Summary of 3 trials to assess causal prophylactic efficacy of ELQ-331 and other prodrugs The upper table segment compares 30 mg/kg prodrug doses; the middle segment, 10 mg/kg prodrug doses; and the lower segment compares 2.5, 5, and 10 mg/kg ELQ-331 doses. Results show more durable protection with ELQ-331 (30 mg/kg, >4 months; 10 mg/kg, >3 months) than other tested prodrugs, and MEC_LAI-C_ between 60-80 nM (see text)

| Treatment group | Time from treatment to sporozoite challenge | Luminescence imaging results |  |  | Closest corresponding [ELQ-300], nM (time, mean±SEM, individual values) |
| --- | --- | --- | --- | --- | --- |
|  |  | Liver stage (2 days after challenge) | Blood stage (5-6 days after challenge) | Mice protected from infection |  |
| Controls | n/a |  |  | 0/2 (0%) | n/a |
| ELQ-331, 30 mg/kg (Trial 1) | 137 d |  |  | 3/3 (100%) | 120 d, 104.9±15.2, 134, 92, 88 |
| ELQ-499 | 137 d |  |  | 2/3 (67%) | 120 d, 72.4±13, 98, 62, 57 |
| ELQ-500 | 137 d |  |  | 1/4 (25%) | 120 d, 65.5±40, 185, 37, 22, 18 |
| ELQ-331, 30 mg/kg (Trial 2) | 48 d |  |  | 4/4 (100%) | 49 d, 213.7±25.4, 270, 229, 208, 148 |
| Controls | n/a |  |  | 0/4 (0%) | n/a |
| ELQ-331, 10 mg/kg | 77 d |  |  | 3/4 (75%)*<br>1/4 (25%) by microscopy | 77 d, 67.5±11.4, 92, 75, 66, 37 |
| Controls | n/a |  |  | 0/4 (0%) | n/a |
| ELQ-331, 10 mg/kg | 29 d |  |  | 4/4 (100%)<br>4/4 (100%) by microscopy | 27 d,<br>95.8±3.7,<br>104, 100, 90, 89 |
| ELQ-331, 5 mg/kg | 29 d |  |  | 4/4 (100%)*<br>2/4 (50%) by microscopy | 27 d,<br>50.3±5.7,<br>61, 58, 46, 36 |

\* Whole animal imaging was the *a priori* measure of efficacy, however, blood smear revealed occult infection – see text

In Trial 3, snapshot PK determinations 30 and 70 days after injection of 10 mg/kg showed that sustained [ELQ-300] exposure was greater after ELQ-331 than after injection with long-chain variants ELQ-494 or ELQ-495 (see Figure 4, table insert). ELQ-331 thus remained our lead, and sporozoite challenge to these mice was done 77 days after injection (see Table 3, middle segment). Whole-body imaging of the mice treated with 10 mg/kg ELQ-331, showed no evident liver stage infection, and a faint signal indicating blood stage infection in 1 of 4 animals. Liver stage infection was evident in only 3 of 4 control mice but blood stage infection subsequently occurred in all 4, illustrating that the sensitivity of imaging is not 100%. Although whole-body luminescence imaging is our primary tool and *a priori* measure to determine outcome after sporozoite challenge, we sought to gain added sensitivity by examining blood smears of the four ELQ-331-treated mice immediately after imaging. Parasitemia was <0.1% in the mouse that was faintly-positive by imaging, <0.01% in 2 mice, and one mouse was parasite-free. On the day of imaging, [ELQ-300] values measured in these mice were 92, 75, 66, and 37 nM, suggesting 80 nM as an initial estimate of MEC_LAI-C_.

**Figure 4:**
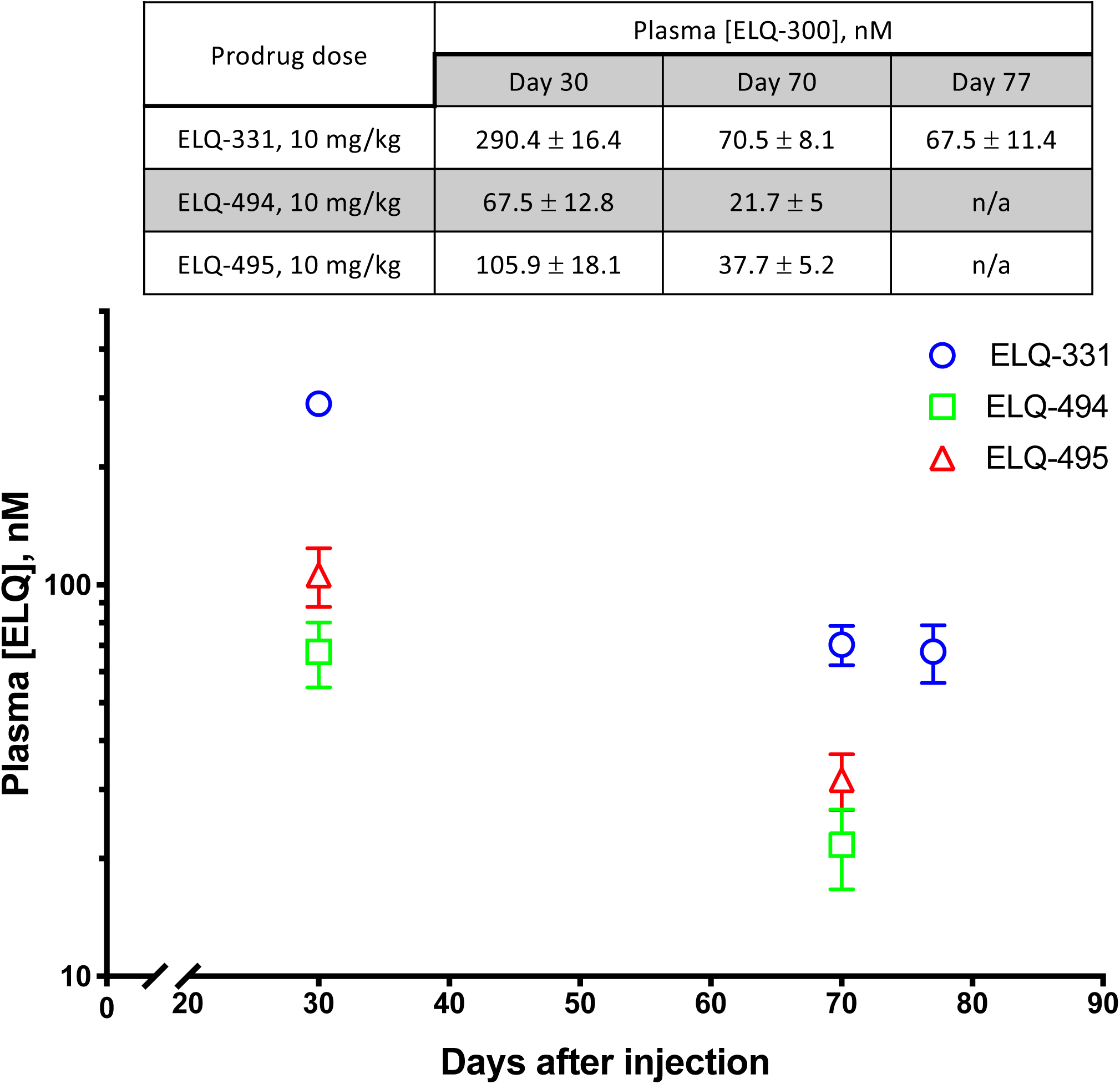
Snapshot pharmacokinetics of ELQ-300 during Trial 3 Snapshot plasma [ELQ-300] values from Trial 3, comparing IM injections of 10 mg/kg ELQ-331 with two other ELQ-300 prodrugs. The log-linear graphical representation of [ELQ-300] vs. time and summary table both display mean [ELQ-300] ± SEM.

To begin to assess the relationship between dose and [ELQ-300] within the first month after ELQ-331 treatment, and to further explore the MEC_LAI-C_, Trial 4 was initiated to compare 10, 5, and 2.5 mg/kg doses of ELQ-331, with 8 mice in each treatment and control group. Snapshot [ELQ-300] determinations 14 days after dosing in mice from each treatment group (N=4) revealed plasma [ELQ-300] values of 183±29, 89±10, and 41±7 nM, respectively (see Figure 5, table insert). Measurement of [ELQ-300] was repeated 27 days after dosing in the other 4 mice from each group and, excepting the 2.5 mg/kg group with earlier sub-effective [ELQ-300], these same mice then underwent sporozoite challenge two days later (see Table 3, lower segment). Mice treated with either 5 or 10 mg/kg of ELQ-331 showed no liver or blood stage infection evident by imaging 2 or 5 days after inoculation, whereas all controls were positive at both timepoints. Blood smears from the mice in the 5 and 10 mg/kg groups were made 7 days after sporozoite challenge to rule out low-grade infection below the level of detection by imaging. No parasites were found among the mice treated with 10 mg/kg or in two of the mice treated with 5 mg/kg, but smears revealed 1% parasitemia in the other two animals. Weekly blood smears remained parasite-free for 28 additional days in the uninfected mice. In the 5 mg/kg treatment group [ELQ-300] values were 61, 58, 46, and 36nM, suggesting that in this trial, an estimate of 60 nM might be appropriate for the MEC_LAI-C_.

**Figure 5:**
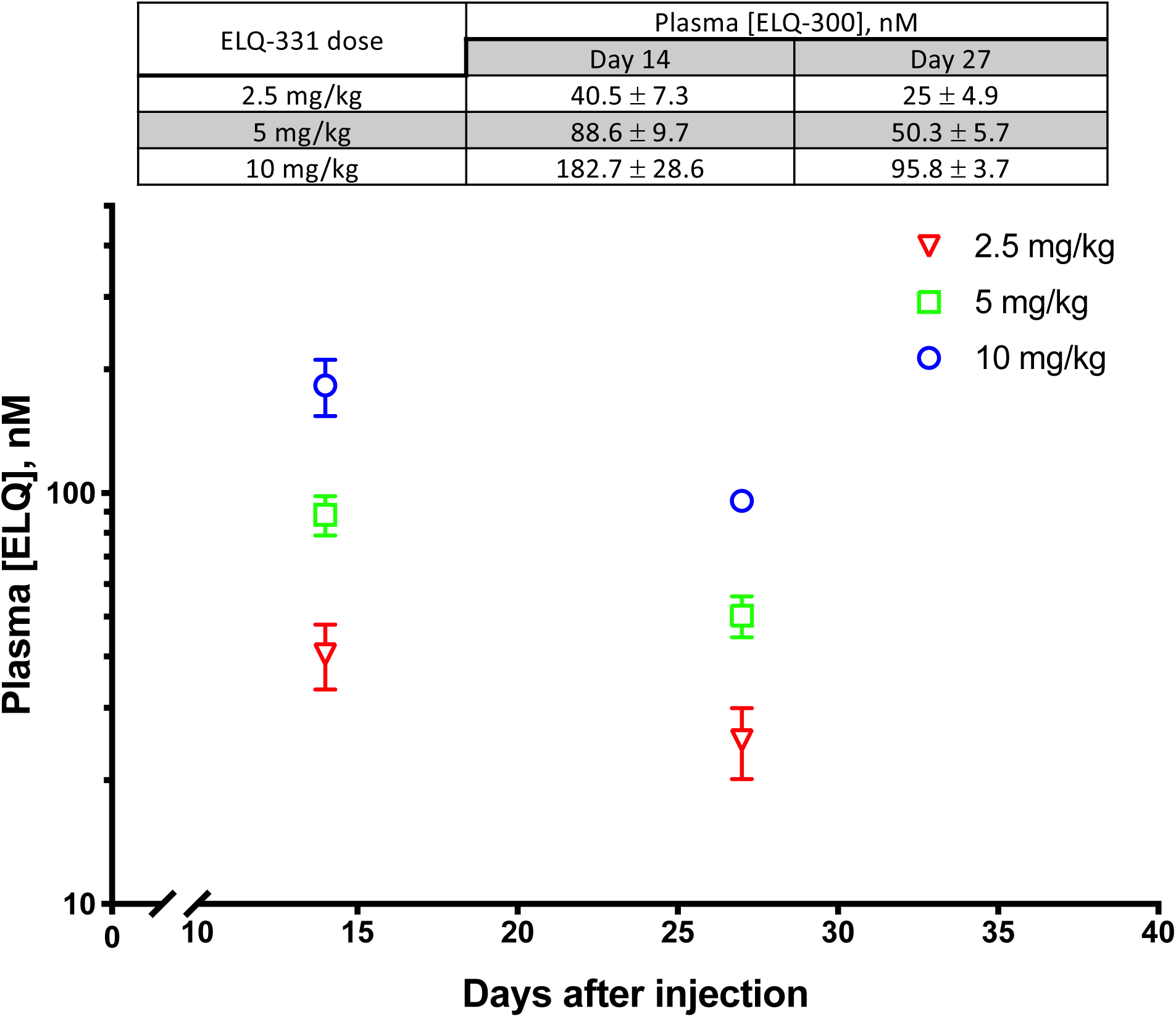
Snapshot pharmacokinetics of ELQ-300 during Trial 4 Snapshot plasma [ELQ-300] values from Trial 4, comparing IM injections of 2.5, 5, and 10 mg/kg ELQ-331. The log-linear graphical representation of [ELQ-300] vs. time and summary table both display mean [ELQ-300] ± SEM.

This value is substantially lower than the 80 nM value described above, and the source of this difference is unclear, but we suspect that it may be the result of differences in the effective inoculum. In the latter experiment, there was a much weaker liver stage infection signal in control mice, suggesting that the actual delivered inoculum of viable sporozoites was smaller, possibly the result of differences in salivary gland transport time and sample processing. The latter experiment also used CF-1 mice, while the mice in earlier trials were CFW (CF-1 mice were temporarily unavailable from the supplier), although this seems unlikely to account for either reduced infectivity of the Py-GFP-luc sporozoites, or enhanced potency of ELQ-300. Since it is unclear how experimental Py-GFP-luc inoculum size relates to predicting efficacy against natural infection in humans, it seems prudent to consider 80 nM as a conservative target MEC_LAI-C_ for LAI-C studies using the Py-GFP-luc sporozoite model in mice.

We did not design trials to assess MEC separately for causal and suppressive prophylaxis, but we believe these MEC_LAI-C_ values to reflect true causal prophylaxis, and not suppression of early blood stage infection. Per protocol, mice with any evident infection after sporozoite challenge were euthanized, so the only opportunity to assess possible suppressive efficacy was in the mice with normal imaging results and no parasites on initial blood smear. As noted above, 2 of those mice developed approximately 1% parasitemia within 2 days, a rapid growth rate suggesting that [ELQ-300] was well below the effective threshold against blood stage infection. Mice with only slightly higher [ELQ-300] were completely protected, making it more likely that these values approached the threshold for causal prophylactic efficacy instead. This is consistent with study findings after IM atovaquone, in which concentrations that protected against sporozoites failed to protect against blood stage challenge (7).

## Discussion

These results show that a single IM injection of 30 mg/kg ELQ-331 provides more than 4 months of causal prophylaxis against malaria in mice and has a distinctive PK profile which contributes to its durable efficacy. These results were achieved using unaltered ELQ-331 dissolved in sesame oil and 1.2% benzyl alcohol (as a preservative), without either vehicle optimization or formulation engineering, suggesting that further enhancements are likely possible. Translation of these results to predictions of efficacy in humans is speculative, but standard allometric scaling (26) provides reason for optimism. If the in vivo ELQ-300 MEC_LAI_ against species causing malaria in humans is similar to that determined in mice, allometric scaling would predict that the duration of effect of a given ELQ-331 dose would be several-fold longer in humans than in mice, or, conversely, that a far lower dose would provide the same duration of effect. Taken together, these observations suggest a very high likelihood that ELQ-331 can feasibly provide the 3 months of protection currently called for in the MMV’s TPP for LAI-C against malaria.

These results are also impressive in comparison to the scant available results from other drugs tested for LAI-C. In two studies of engineered drug particles injected IM, mice were protected against *P. berghei* sporozoite challenge for 56 days by a microsuspension of decoquinate (120 mg/kg) in peanut oil (8), and for 28 days by atovaquone nanoparticles (200 mg/kg) (7). In unpublished studies of IM prodrugs of atovaquone and ELQ-300, apparent MEC_LAI-c_ values for atovaquone and ELQ-300 were established based on protection against sporozoite challenge 14 days after injection, and duration of efficacy was then predicted based on how long the MEC_LAI-C_ was exceeded (Presentation by Chatterjee, A.K., *TCP4: Intra-muscular injections for malaria chemoprotection*, in *Symposium: What kind of molecules are needed to control and eradicate malaria? ASTMH 66th Annual Meeting*. 2017: Baltimore, MD). No later sporozoite challenges were reported, but after 30 mg/kg injections, MEC_LAI-C_ was exceeded for more than 13 weeks in dogs after the ELQ-300 prodrug (mCBK069), and in rats after the ATV prodrug (mCBK068). Given allometric scaling predictions (clearance in mice < rats << dogs), the ELQ-331 performance demonstrated here appears to surpass that of other agents for which reports are available.

Although sesame oil is used as a vehicle for other LAI drugs, such as haloperidol (5), it is unlikely to match the initial specifications set out by the MMV. The TPP for antimalarial LAI-C calls for a small volume injection, ideally 1 mL or less in adults (< 2 mL is acceptable) and 0.5 mL or less in infants, and that it be injectable through a 27-gauge needle (9). We find ELQ-331 in refined sesame oil to be soluble to 83 mg/mL (see Additional File 2). Given that 30 mg/kg of ELQ-331 yields 4-5 months of protection in mice, and allometric scaling would suggest 2.5 mg/kg as an equivalent dose in humans, it may or may not prove difficult to deliver the dose needed for 3 months of protection for a large adult in 1 mL. Due to viscosity, delivery through a very fine needle is problematic for any conventional oil. Oily vehicles enhance the solubility of lipophilic drugs such as ELQ-331 and serve to slow release from the injection site, and it is unclear how difficult it will be to achieve success with a low viscosity vehicle. ELQ-331 is soluble in some low viscosity vehicles, such as Miglyol 840 (see Additonal File 2), and the results of solubility testing in several vehicles provide a starting point for optimization. This suggests that adequate solubility will be achievable, but it will also be necessary for the vehicle to have a desirable drug release profile, local and systemic tolerability, and to be approved for injection use. Viscosity, surface area, and composition of the vehicle all impact the time course of drug release from the injection site (24), so substantial testing may be needed to find the ideal formulation.

Given the many advances in dosing formulation technology, there is little doubt that an ideal vehicle is possible, but for large-scale use in malaria-endemic regions it will be essential to keep costs low and avoid complex vehicle formulations, if possible. If finding an ideal, cost-effective vehicle formulation proves difficult, then it is worth discussing whether inexpensive oil vehicles, such as sesame oil, deserve consideration for LAI-C dosing. Certainly, use of the smallest bore needles will not be possible with oils, but that is a lesser concern. Although the least possible injection discomfort is always desirable, immunizations are routinely injected in infants via 23-25 g needles, so if it were ultimately not possible to use a 27 g needle that would likely still be acceptable. Another concern expressed with regard to sesame oil is allergy risk, but it is uncertain whether such a risk actually exists. There are clearly cases of eosinophilic pneumonitis after repeated daily injections of sesame oil (27), but this results from delayed hypersensitivity to significant amounts of absorbed oil and not immediate hypersensitivity-mediated acute allergic reaction. The antigens in sesame oil capable of causing dangerous acute allergic reactions are present in crude sesame oil, but not in oil refined for pharmaceutical use and most injectable pharmaceutical products in sesame oil do not include allergy risk as a contraindication or potential adverse effect in product information (28). We were unable to find any documented cases in the literature of dangerous acute allergic reaction to pharmaceutical sesame oil, and would argue that it should not be ruled out on the basis of allergy risk. Beyond conventional IM injection, LAI-C is also an excellent candidate application for other delivery systems such as subcutaneous implant (29) and in situ gelling (30) technologies, providing numerous additional options for solubilization and delivery.

To account for the sustained efficacy of ELQ-331, it is useful to first consider its resulting overall [ELQ-300] PK profile. During the initial downslope phase (days 7-35), absorption from the injection site, distribution to and from tissue compartments, and elimination are likely occurring. ELQ-300 is highly metabolically stable, so it is presumed that simple first order processes lead to elimination in feces and urine, driven by mass action, and as a result, ELQ-300 elimination should follow a consistent, concentration-dependent curve. Therefore, the inflection to a decreased slope of the curve (days 45-90) must result from some combination of continued drug absorption, decreased distribution from blood to tissue, and increased reabsorption from tissue to blood. While some drug absorption could still be occurring at that time, for it to be the predominant cause of this plateau phase would require that fairly constant and substantial absorption be ongoing for 3 months or more; this is inconsistent with studies of the persistence of oil and drug at intramuscular injection sites in mice (25). Consistent with our compartmental PK modeling, it is more likely that redistribution from other tissue compartments to blood is predominant over the flat portion of the curve, and that the subsequent increased curve slope, evident after 90 days, results as redistribution from tissue decreases and elimination becomes gradually more predominant. This final apparent change in slope is not accounted for in our standard compartmental modeling equations, and it will be interesting to see in future studies if it is a reproducible PK feature or merely an artifact due to our sampling protocol.

A full, accurate, and predictive model of IM ELQ-331 pharmacokinetics will be complex since it must include the rate of release of ELQ-331 from the injected oil depot; conversion to ELQ-300 by esterases at the injection site, in lymph, blood, and in liver and other organs; absorption into blood; extensive protein binding in blood; distribution from blood to tissue; redistribution from tissue to blood; and elimination, with each component involving its own set of numerous variables (24, 31). As a result, the absolute values of parameters estimated from our simple compartmental PK analyses are unlikely to represent physiologic values in vivo, but the findings are powerful, nonetheless. Fundamentally important is the demonstration that prolonged redistribution of ELQ-300 from tissue sites is critical to the sustained elevation of [ELQ-300] seen here. While the slow release of highly-lipophilic ELQ-331 from oil and the long-elimination T_1/2_ of ELQ-300 certainly contribute to a long duration of effect, these are properties shared by other antimalarials, whereas the capacity of a large, slow-exchange tissue compartment to impact blood drug concentration and efficacy months after dosing is highly unusual, and distinguishes ELQ-300 prodrugs such as ELQ-331 from other candidates reported thus far.

If the ideal PK profile for LAI-C is to rapidly achieve and maintain effective but non-toxic concentrations of the active drug, in most cases this would be achieved by balancing elimination with sustained and controlled release from the injection site throughout the period of risk. For medications delivered by LAI, drug concentrations are typically most dependent on controlled and sustained release from the site of delivery, optimized by choice of vehicle, vehicle mixtures, benzyl alcohol or other excipient content, drug particle engineering, or other injection parameters (24, 29). This is evident in unpublished data regarding atovaquone prodrugs (32), in which day 30 [atovaquone] values are highly dependent on formulation. In contrast, our results suggest that maintaining sustained and effective ELQ-300 concentrations after IM ELQ-331 is most closely related to loading and gradual unloading of ELQ-300, to and from a repository tissue depot. While the pattern of ELQ-331 release from the injection site is undoubtedly important in this process, controlled release apparently does not need to be sustained to sustain durable protection because of the effect of the secondary depot. Because this PK profile is a distinguishing feature of ELQ-331, it is important to consider its possible disadvantages and advantages in the setting of LAI-C.

It will be important to identify the tissues and organs comprising this large-capacity, slow exchange compartment to understand its impact on dosing considerations. If distribution is highly concentrated in specific organs (e.g., liver), between-subject variability on the basis body mass index (BMI) obesity would be less likely, whereas variations in BMI might lead to unwanted variations in response if distribution is predominantly to general adipose tissue. Further study will allow development of the physiology-based PK models needed to address these questions.

Extensive and sustained tissue stores of ELQ-300 could, theoretically, also prevent rapid termination of exposure in case of toxicity, particularly if the site of toxicity were an organ in which the drug is highly concentrated. After typical IM injection, this risk would be similar for any drug, but since removable drug delivery systems (e.g., implants, removable gels) are a consideration for LAI-C as a safeguard against toxicity, in that context a rich deposit of tissue-bound drug would be a comparative disadvantage. At this time, there are no known safety concerns with ELQ-331, but with large required drug payloads, sustained drug exposure, and planned use in remote settings with little safety surveillance capacity, the safety of any drug used for LAI-C must be viewed skeptically.

During oral toxicity testing of ELQ-331, rats received a single dose of 15, 30, 100, 300, or 1000 mg/kg, or 7 daily doses of 5, 50, or 300 mg/kg (see Additional File 1). The lowest single dose with any evident toxicity was 300 mg/kg, which resulted in an ELQ-300 C_max_ = 65-79 μM, and after lower doses, concentrations well above 27-44 μM caused no apparent adverse effect. Seven daily doses of 5 mg/kg (35 mg/kg cumulative) were tolerated without evidence of toxicity; [ELQ-300] values after the last dose showed C_max_ = 13-17 μM, and indicated that supra-μM concentrations had been maintained throughout the week. While neither single nor 7 day dosing mirrors exposure after LAI-C, the very high [ELQ-300] values associated with toxicity and the absence of toxicity after either a single 100 mg/kg or cumulative 35 mg/kg over 7 days are reassuring that the 4-month duration of protection reported here was achieved safely (C_max_ = < 5 μM).

If safety is ultimately established, then the ELQ-331 PK profile could be highly advantageous. By exploiting intrinsic PK properties to maintain effective ELQ-300 concentrations, a shorter period of tightly-controlled release might be needed, lessening the requirement for extensive formulation engineering. In fact, if “loading” this compartment is sufficient to sustain concentrations, then this might be achievable by many drug delivery routes, including short-term repeated oral dosing. Essentially, this property adds another basis for extended efficacy, regardless of dosing strategy, and suggests that duration of effect for far longer than 3 months may be more easily achievable with ELQ-300 prodrugs than with atovaquone derivatives or other candidates.

This is advantageous beyond simply meeting the initial 3-month duration-of-effect target. To avoid conditions favorable to generation of drug resistance during proposed combination therapy, it is important that partner drugs are matched such that neither component falls below the MEC_LAI_ during the period of risk. Achieving such a match is challenging, particularly if both components require significant engineering to meet duration-of-effect requirements. If ELQ-331 requires little or no engineering to achieve adequate extended activity, then adjusting duration of effect to match any partner drug should be simplified, perhaps only requiring dose adjustment. Alternatively, in settings with a defined finite period of exposure risk, ELQ-331 might facilitate an acceptable PK mismatch in which the effectiveness of a partner drug extends to the end of the risk period, but ELQ-331 effect continues beyond that time. Finally, more reliable sustained efficacy may support consideration of ELQ-331 for use without a partner.

Avoidance of monotherapy is based on the concern that a large number of parasites in the presence of sub-effective drug concentrations promotes the generation or selection of resistant parasites. Clearly a risk during blood stage infection, the same risk is theoretically possible during late liver stage infection when infected hepatocytes house many thousands of developing merozoites. In contrast, an effective causal prophylactic that prevents liver schizont development and survival faces less than a hundred non-replicating parasites, making the risk of resistance generation exceedingly low if not zero. In the absence of pre-existing resistance, monotherapy LAI-C is thus a logical consideration if an effective drug concentration can safely and reliably be maintained beyond the end of exposure risk. This argument applies only to true causal prophylaxis, so it is important to distinguish whether efficacy is solely on that basis, or whether results are confounded by suppression of blood stage growth. ELQ-300 retains in vitro potency against *P. falciparum* clinical isolates resistant to quinolines, anti-folates, artemisinin, and atovaquone, among others, with no evidence of existing resistance. Generation of ELQ-300 resistance in vitro is several orders of magnitude more difficult than with atovaquone, and like the case of atovaquone, there is evidence of diminished transmissibility of ELQ-300 resistant parasites (personal communication, G. McFadden). Given its multistage efficacy, potency, PK profile, and activity against multidrug resistant parasites, ELQ-331 seems an ideal candidate for consideration of study as single-drug LAI-C, in addition to its less controversial proposed role in 2-component LAI-C.

To meet shelf-life requirements for LAI-C, ELQ-331 would need to undergo full stability testing in whichever formulation is ultimately selected for LAI-C, but existing results are encouraging. Testing for 3 months at 40° C and 75% humidity showed ELQ-331 in a spray-dried dispersion formulation to be stable (manuscript submitted), and while no long-term testing has been done, it is stable in the short-term after heating in sesame oil. Regarding the affordability of ELQ-331 as a component of LAI-C, there are no obstacles to scalable, feasible, and cost-acceptable ELQ-331 synthesis, and only the need for an unexpectedly complex and expensive final formulation could raise costs to a concerning level.

The above findings suggest that ELQ-331 is an excellent candidate for use in LAI-C against malaria and, that among current tier 2 candidates, its characteristics are distinctly and perhaps uniquely suited to that purpose, making it an appropriate prototype for study and comparison. Further studies are warranted, including expanded toxicology studies of ELQ-331 relevant to LAI-C, consideration of appropriate partner drugs, study of factors relevant to the consideration of monotherapy, exploration of other vehicle formulations for IM injection, study of other delivery systems (e.g., implant, in situ gelling polymer, oral), and continued comparison with other prodrugs.

## Summary

A single IM injection of ELQ-331 in sesame oil in mice provided causal malaria prophylaxis for more than 4 months. Given allometric scaling predictions and available engineering and formulation technologies, these results clearly support the feasibility of ELQ-331 to achieve duration of protection against malaria for at least 3 months in humans. Furthermore, the distinct PK profile of ELQ-300 after ELQ-331 treatment may facilitate flexible delivery and formulation options and may also enable protection for far longer than 3 months. Particularly in light of the favorable pharmacodynamic profile of ELQ-300, ELQ-331 warrants consideration as a leading tier 2 prototype for LAI-C.

## Abbreviations

BMI: Body mass index
C_max_: Maximum concentration
[ELQ-300], [ELQ-331]: Concentration of ELQ-300, concentration of ELQ-331
G-6-PD: Glucose-6-phosphate dehydrogenase
IS: Internal standard
IM: Intramuscular
LAI-C: Long-acting injectable chemoprotection
MEC: Minimum effective concentration
MEC_LAI-C_: Minimum effective concentration for chemoprotection after long-acting injection
MMV: Medicines for Malaria Venture
MRM: Multiple reaction monitoring
PK: Pharmacokinetic
TPP: Target product profile

## Declarations

### Authors’ contributions

R. A. D., L. F., S. P., R. W. W., A. K., A. N., and J. X. K. made possible the design, synthesis, purification, and analysis of ELQ-300 and its prodrugs. For Py-GFP-luc sporozoite challenge experiments, S. W., T. M., B. K W., and S. H. I. K. each were essential participants in the experimental design; parasite supply; and sporozoite production, harvest, and processing. Y. L. and D. J. H. performed drug dosing, *P. yoelii*_*K*_ and Py-GFP-luc inoculation, and blood smears. M. J. S. and Y. L. performed animal monitoring, sampling for pharmacokinetic analysis, and initial sample processing. I. B. performed the preparation, execution, and image processing for whole-body imaging. Analyses of [ELQ-300] and [ELQ-331] were performed by L. A. B. and D. R. K. Pharmacokinetic data analysis was performed by M. Y. M., S. P., and M. J. S. Research design, analysis, and interpretation of results, was coordinated and led by M. K. R. and M. J. S., with all authors providing essential input. The manuscript was written by M. J. S., with several authors contributing.

#### Acknowledgements

We gratefully acknowledge the guidance and support of Jeremy Burrows and the MMV, for making possible the foundational discovery and development research behind ELQ-300. As part of that work, Sue Charman and Karen White of the Centre for Drug Candidate Optimisation at Monash University provided invaluable PK analysis of ELQ-300 by oral and parenteral routes in multiple animal species. Their methods and results were essential background for this project.

### Availability of data and materials

The datasets used and analyzed during the current study are included in this report, in the Additional Files, or are available from the corresponding author on reasonable request.

### Consent for publication

Not applicable.

### Ethics approval and consent to participate

All animal studies were performed under Portland Veterans Affairs Medical Center Institutional Animal Care and use Committeee approved protocols. All animal use, care, and handling were performed in accordance with the current “Guide for the Care and Use of Laboratory Animals” (8th Edition, 2011).

### Disclaimer

The views expressed in this article are those of the authors and do not necessarily reflect the position or policy of the Department of Veterans Affairs or the United States government.

## Supplementary material

*Additional File 1:* Toxicity and toxicokinetic study of ELQ-331 and ELQ-300 following a 7-day dose administration in Sprague Dawley rats

*Additional File 2:* Assessment of a method to limit ex vivo conversion of ELQ-331 to ELQ-300; Generalized method for estimation of compound solubility in sesame oil; and ELQ-331 solubility in several solubilizing excipients

*Additional File 3:* Pharmacokinetic modeling

*Additional File s:* Pharmacokinetic raw data

