## Additional File 1 for "ELQ-331 as a prototype for extremely durable chemoprotection against malaria"

**Additional File 1: *Toxicity and toxicokinetic study of ELQ-331 and ELQ-300 following a 7-day dose administration in Sprague Dawley rats*** *–* (SRI Study Number: M380-18, SRI International, Menlo Park, CA)

The objective of this study was to determine 1) the maximum tolerated dose (MTD) after a single oral dose of the prodrug ELQ 331; and 2) toxicity and toxicokinetics (TK) after a 7-day repeat dose of ELQ 331 in male and female Sprague Dawley rats following an oral (po) administration.

The study was conducted in two phases. The maximum tolerated dose (MTD) phase (Phase A) was performed by administration of a single oral dose of the prodrug ELQ-331 at 15, 30, 100, 300 or 1000 mg/kg (Groups 2-6, respectively) to male and female Sprague Dawley rats (3/group/sex). A control group (Group 1) was given a single oral dose of vehicle (37.5% PEG 400, 37.5% Tween-20 and 25.0% Capmul MCM NF). Endpoints of toxicity including clinical observations and body weights were determined. Clinical findings were observed in Group 5 (300 mg/kg) animals that included drooling and slight hypoactivity. Clinical findings observed in Group 6 (1000 mg/kg) included drooling, dyspnea, sneezing, hypoactivity, squinting and skin and fur discoloration and fecal staining, rales, ruffled fur and soft stool. Body weight loss was observed throughout the study only in animals at 300 and 1000 mg/kg.

Based on these findings, the dose levels for the repeat dose and TK phase (Phase B) of the study were selected to be 5, 50 and 300 mg/kg (Groups 8-10, respectively) for daily administration for 7 days to male and female Sprague Dawley rats. The following parameters were evaluated: mortality and morbidity, clinical observations, body weights, clinical pathology, organ weights, macroscopic and microscopic evaluations.

In Phase B, clinical findings including dehydration and ruffled fur were observed at 50 mg/kg which appeared to be test article-related. In Group 10 (300 mg/kg), hypoactivity, ruffled fur, dehydration and soft stool were observed in all animals after repeat dose administration. Two females in Group 10 appeared thin on Days 5-8. These observations are considered to be test article related. Other clinical findings were observed throughout the study beginning on Day 2 including slight to extreme fecal stain, moderate urine stain, shoveling, slight drooling, anus and muzzle skin and fur discolored, and wet anus covered with fecal and urine stain. Suppression of body weight gain and weight loss were observed throughout the study in the 50 and 300 mg/kg dose groups. At the scheduled necropsy on Day 8, several white blood cell (WBC) parameters were changed in both groups, including increases in neutrophils, lymphocytes and monocytes and decreases, in males only, in reticulocytes. Animals of these groups also showed increases in aspartate aminotransferase (AST) and alanine aminotransferase (ALT). Dose dependent decreases in thymus weight, thymus-to-body weight and thymus-to-brain weight were observed in both males and females at the 50 and 300 mg/kg dose levels. The weight changes in thymus and small thymus in macroscopic finding are consistent with the histopathological findings. At the 300 mg/kg dose level, increases were observed in liver and spleen weights. These changes appeared to be test article-related and are consistent with the histopathological findings described below.

Test article-related histopathologic findings associated with ELQ-331 administration included increased hepatocellular mitosis, hepatic karyomegaly, decreased lymphoid cellularity of the thymus and spleen, increased lymphoid apoptosis of the thymus, gastric erosion/ulcer, seminal vesicle epithelial vacuolation, kidney vacuolation, and macrophage or macrophage-derived cell accumulations within the spleen, lung, thymus, and liver (Kupffer cells). The observation of foamy/vacuolated macrophages or monocyte-derived cells (Kupffer cells) in various tissues (spleen, liver, lung, and thymus) and tissue vacuolation (seminal vesicle and kidney) may be suggestive of accumulation of phospholipids in this study.

Toxicokinetic data analysis was performed on the plasma concentration data of ELQ-331 and its active metabolite, ELQ-300 (see Table, below). Plasma concentrations of ELQ-300 were much greater than those of ELQ-331, with up to almost 2,000-fold greater AUC_last_ values for ELQ-300 than ELQ-331. Plasma exposure of ELQ-331 in some groups decreased with increasing doses and on Day 7 versus Day 1, suggesting a possible saturation of metabolite formation with dose increments and repeat dose administration with ELQ-331. Correspondingly, the formation of the active metabolite ELQ-300 was less than dose proportional and repeat dose administration of ELQ-331 resulted in reduced formation of ELQ-300 by Day 7. No major trends were noted in sex differences in the toxicokinetics of the ELQ-331 and ELQ-300. For comparison to PK results after LAI-C in mice, note that 1 ng/mL = 2.1 nM.

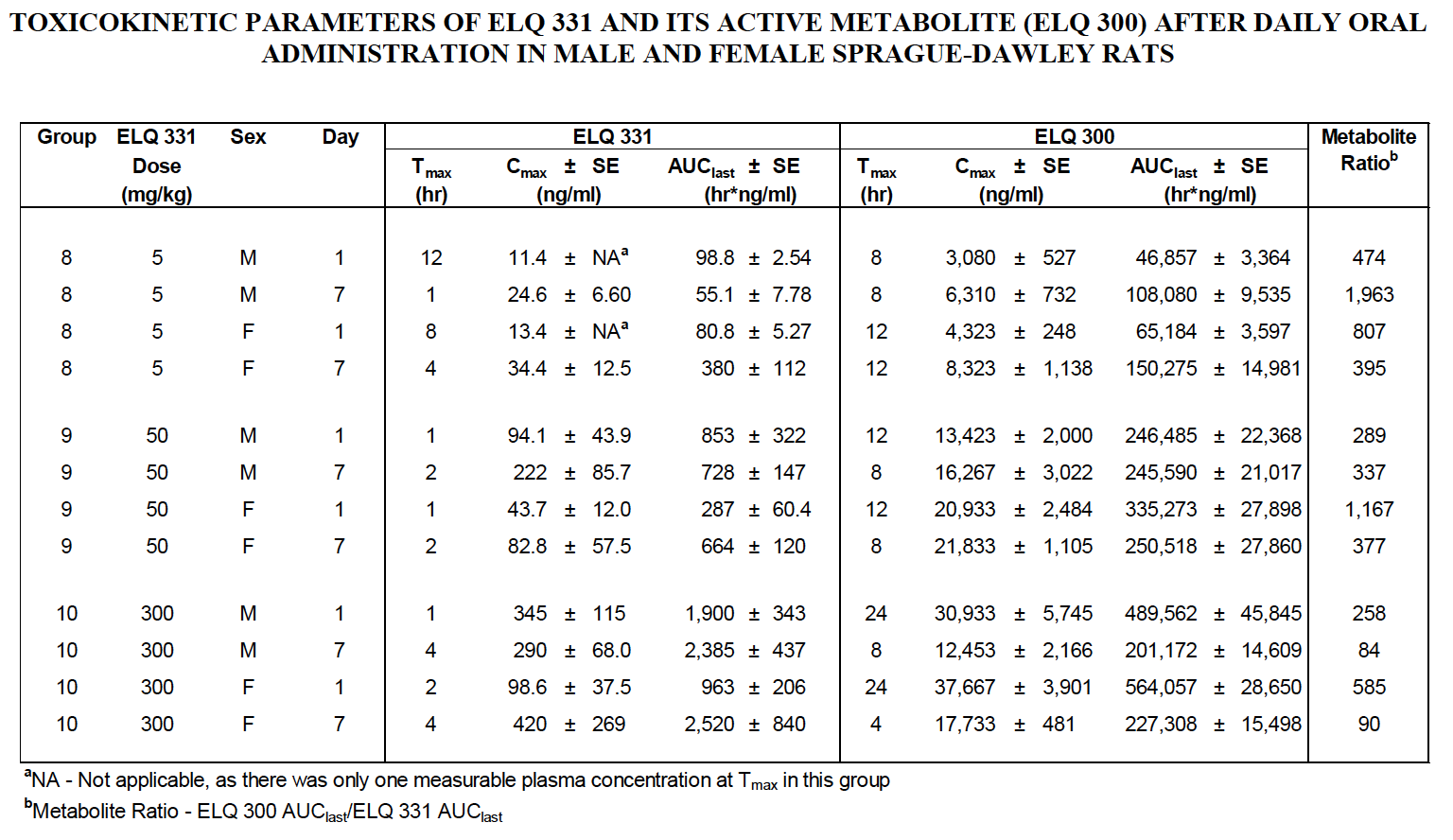
In conclusion, a single oral dose of ELQ-331 to male and female Sprague Dawley rats did not produce toxicity at 15, 30 or 100 mg/kg while clinical signs and body weight loss were observed at 300 and 1000 mg/kg. Repeat dose administration for 7 days was well tolerated at 5 mg/kg, but clinical signs, body weight loss, changes in white blood cells, increased levels of AST and ALT and histopathologic changes were observed at 50 and 300 mg/kg (cumulative 350 and 2100 mg/kg, respectively). Based on these results, the maximum tolerated dose (MTD) of ELQ-331 after 7 days of oral dose administration in Sprague Dawley rats is considered to be approximately 300 mg/kg. While some microscopic findings were observed at 5 mg/kg, these are all considered of negligible toxicologic significance; therefore the no-observed-adverse-effect-level (NOAEL) is considered to be 5 mg/kg when administered orally for 7 consecutive days.
