## Additional File 2 for "ELQ-331 as a prototype for extremely durable chemoprotection against malaria"

1. ***Assessment of a method to limit ex vivo conversion of ELQ-331 to ELQ-300***
2. ***Generalized method for estimation of compound solubility in sesame oil***
3. ***ELQ-331 solubility in several solubilizing excipients***

***Assessment of a method to limit ex vivo conversion of ELQ-331 to ELQ-300****:* Using a heparinized syringe, 1 mL of blood was obtained from a female CFW mouse by cardiac puncture, immediately spiked with 1 µL of 10 mM ELQ-331 in DMSO, and mixed thoroughly. From this sample, 300 µL aliquots were transferred to three microfuge tubes: one at room temperature, one on ice, and one containing 34.2 µL of 500 mM sodium fluoride (NaF) on ice. Tubes were left under these conditions for one hour, then centrifuged at high speed for one minute. Aliquots of supernatant plasma were removed from each tube, and immediately frozen at -80° C until determination of plasma ELQ-300 concentration ([ELQ-300]). The results summarized below indicate that ex vivo hydrolysis of ELQ-331 to ELQ-300 is effectively limited by the combination of cold and NaF, and has little or no impact on plasma [ELQ-331] and [ELQ-300] values determined during our PK analysis.

| Condition tested | Initial [ELQ-331]_calculated_ | Final [ELQ-300]_measured_ | Estimated ex vivo hydrolysis of ELQ-331 |
| --- | --- | --- | --- |
| None | 10 µM | 727 nM | 7.3% |
| Cold | 10 µM | 387 nM | 3.9% |
| NaF, Cold | 10 µM | 194 nM | 1.9% |

***Generalized method for estimation of compound solubility in sesame oil*:** Sesame oil (10µl) was dispensed into a glass vial containing 10 mg of the appropriate ELQ-300 prodrug. The mixture was vortexed and warmed (using a heat gun) to aid dissolution. The solubility of the compound was judged after the sesame oil mixture cooled to room temperature. If the compound was not completely dissolved, additional sesame oil was added in low-volume increments. Importantly, complete dissolution was assured by confirming by visual inspection that the compound remained in solution for at least 72 hours at room temperature.

***ELQ-331 solubility in several solubilizing excipients*** *-* Dissolution of ELQ-331 powder in several known solubilizers (see table, below) was attempted, using warming and mixing only to aid solubilization, and visual inspection with magnification to assess results. The highest concentration tested was 250 mg/mL. If the powder failed to go into solution completely, aliquots of the solubilizer were added in various proportions, and solubility was reassessed. This process was repeated until either a clear solution resulted or until solubility was found to be < 50 mg/mL. Clear solutions were left at room temperature for at least 24 hours and re-assessed, and only those remaining clear were included in determining the values listed below.

| **Vehicle** | **Solubility (mg/ml)** |
| --- | --- |
| Capryol 90 | >250 |
| Glyceryl Tributyrate | >250 |
| Isopropyl Myristate | 63 |
| Labrasol | 125 |
| Miglyol 812 | 83 |
| Miglyol 840 | 200 |
| Olive Oil, refined (Gattefossé) | <50 |
| Sesame Oil, refined (Gattefossé) | 83 |
| Sesame Oil, unrefined | 83 |
| Soybean Oil, refined (Gattefossé) | <50 |
| Transcutol | 167 |
