## Additional File 3 for "ELQ-331 as a prototype for extremely durable chemoprotection against malaria"

**Additional File 3: Pharmacokinetic modeling**

The dataset used for PK modelling included the merged plasma [ELQ-300] values from Trial 1 and Trial 2 (30 mg/kg ELQ-331), with and without editing of values at early timepoints from Trial 2. As discussed in the text, several of the values measured at the very early timepoints after injection displayed large variability, and were not biologically plausible as representing the true pharmacokinetic profile after IM injection. Because the early drug absorption profile is a critical element of modeling, we sought to include and exclude values at early timepoints on the basis of their plausibility as predictive indicators of actual absorption after IM injection ELQ-300 in oil. All values were included in the [ELQ-300] calculations represented in the text and in the raw data (see Additional File 4); these editing adjustments were for PK modeling only.

When considering values to exclude, we used the timepoints at 24 and 36 hours as reference points, since they showed little variability in [ELQ-330] between mice (see main text, Figure 2, insert 1), and it was most plausible that the uniformity at these points indicated the absence of confounding extra-muscular dose deposition. All values at the 24 and 36 hour timepoints were included, and earlier values were assessed in relation. Regarding only time points < 24 hours after injection, we also assumed that if there were both very high and very low [ELQ-300] values from mice within a group, then the low value was likely representative of more complete intramuscular deposition. Using these assumptions, we excluded from modeling consideration some values at the 6, 12, and 18-hour timepoints only in an edited dataset (“Edited”). These data exclusions were made prior to any PK analysis, and both the “Edited” and complete (“All”) datasets were then evaluated for PK profile (see Additional File 4 for complete and modified datasets).

Non-compartmental analysis was done first to determine summary PK values (see Additional File 3, Table 1), and to provide initial parameter estimates for compartmental modeling. Datasets were

then run through 1, 2, and 3-compartment models using SimBiology. Lacking the ability to track and analyze the multiple processes occurring between injection of ELQ-331 and the appearance of ELQ-300 in blood, we chose to treat absorption as a first order process into the central (blood) compartment, speculating that this would better model actual absorption than either a zero order or bolus model. Mean [ELQ-300] values at each time point were used as the dependent variable. Better fit was achieved with the multi-compartmental equations (see Additional File 3, Table 2), with the 2-compartment model slightly better than the 3-compartment model on the basis of fit and error criteria.

**Additional File 3, Table 1: Non-compartmental analysis – Summary values**

| **Parameter, units** | **Estimate** | |
| --- | --- | --- |
| Dataset | "All" | "Edited" |
| Dose, nmol | 1900 | 1900 |
| Points used to calculate terminal T_1/2_ | 5 | 5 |
| Terminal T_1/2_, days | 29.5 | 29.5 |
| Tmax, days | 0.75 | 3 |
| Cmax, nM | 3752.3 | 1614.2 |
| AUC to last timepoint, day*nmol/L | 49215.3 | 46903.1 |
| AUC to ∞ (observed slope), day*nmol/L | 51086.7 | 48774.4 |
| AUC to ∞ (calculated slope), day*nmol/L | 50784.7 | 48472.5 |
| Average [ELQ-300], nM | 294.7 | 280.9 |
| CLss_F, L/day | 0.039 | 0.04 |
| Vz_F, L | 1.64 | 1.73 |

Table legend: Results of non-compartmental analysis and comparison of “All” and “Edited” datasets (see text). AUC = area-under-the-curve, CLss_F = clearance at apparent steady-state, Vz_F = apparent volume of distribution at apparent steady-state

Based on greater plausibility and the initial modeling results showing very poor fit to the “All” dataset, only the “Edited” dataset was further evaluated. The final 2-compartment model is illustrated here (see Additional File 3, Figure 1), along with the corresponding expressions used in the model. The resulting equation (converting The graphical and tabular fit results are presented in the main text (Figure 3) confirming an excellent fit with little difference between observed and predicted [ELQ-300] values and evenly distributed residuals.

**Additional File 3, Table 2: Compartmental analysis – comparison of fit/error values between models**

| **Model - Dataset** | **LogLikelihood** | **AIC** | **BIC** | **DFE** | **RMSE** |
| --- | --- | --- | --- | --- | --- |
| 1-compartment - ”All” | -139.7 | 392.0 | 396.1 | 22.0 | 581.6 |
| 1-compartment - ”Edited” | -193.0 | 285.3 | 288.8 | 21.0 | 87.1 |
| 2-compartment - ”All” | -190.0 | 390.0 | 396.1 | 20.0 | 540.9 |
| 2-compartment - ”Edited” | -129.6 | 269.1 | 275.0 | 19.0 | 60.1 |
| 3-compartment - ”All” | -193.0 | 400.0 | 408.6 | 18.0 | 642.98 |
| 3-compartment - “Edited” | -128.5 | 270.8 | 279.1 | 17.0 | 60.59 |

Table legend: Comparison of fit and error criteria for 1, 2, and 3-compartment PK modeling of both the “All” and “Edited” datasets, showing best fit with 2-compartment modeling of the “Edited” data. AIC = Akaike Information Criterion, BIC = Bayesian Information Criterion, DFE = degrees of freedom for error, RMSE = root mean square error

**Additional File 3, Figure 1: Diagram and expressions for the selected 2-compartment model**

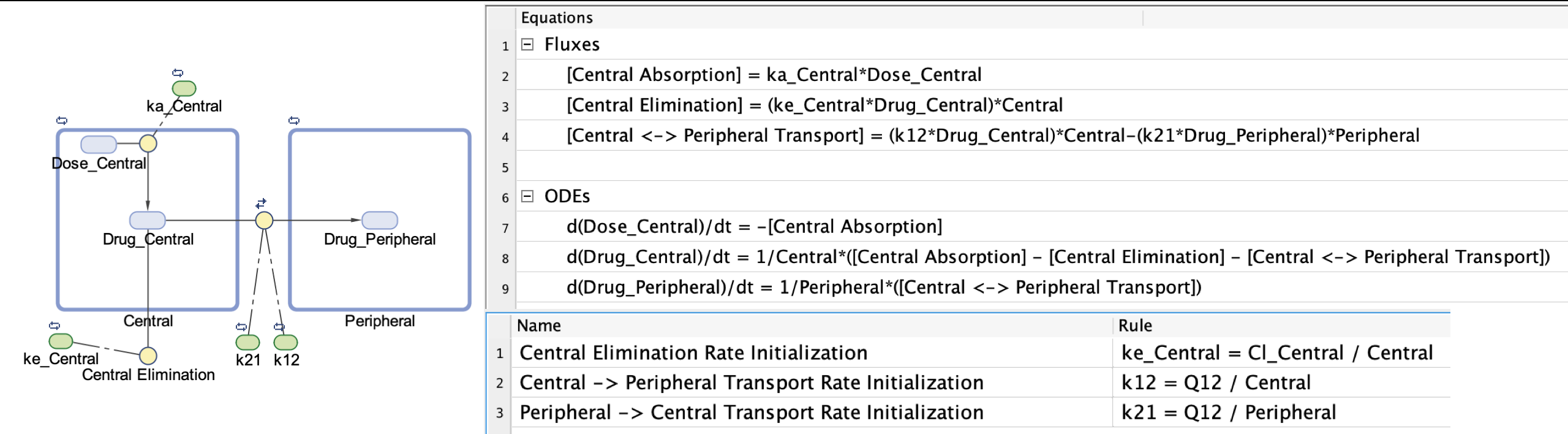
